# Long-B prokaryotic Argonaute systems employ various effectors to confer immunity via abortive infection

**DOI:** 10.1101/2023.03.09.531850

**Authors:** Xinmi Song, Sheng Lei, Shunhang Liu, Yanqiu Liu, Pan Fu, Zhifeng Zeng, Ke Yang, Yu Chen, Ming Li, Qunxin She, Wenyuan Han

**Author notes:** These authors contributed equally to the manuscript (Xinmi Song, Sheng Lei, Shunhang Liu, Yanqiu Liu).

## Abstract

Argonaute proteins (Agos) bind short nucleic acids as guides and are directed by them to recognize target complementary nucleic acids. Prokaryotic Agos (pAgos) are extremely diverse, with potential functions in microbial defense. The functions and mechanisms of a group of full-length yet inactive pAgos, long-B pAgos, remain enigmatic. Here, we show that most long-B pAgos constitute cell suicide systems together with their various associated proteins, including nucleases, Sir2-domain-containing proteins and trans-membrane proteins, respectively. Among them, the long-B pAgo-nuclease system utilizes an RNA-programmed and target-recognition-activated collateral DNA cleavage activity to sense invaders and kill the infected cells. This results in depletion of the invading plasmid from the cell population. Together, our data indicate that the long-B pAgo systems induce cell death with various effector proteins after recognition of invading nucleic acids, corresponding to an immune response via abortive infection.

## Introduction

Argonaute proteins (Agos) are important defense elements in both eukaryotes and prokaryotes ^1^. Eukaryotic Agos (eAgos) provide immunity against viruses and transposons by RNA silencing ^2,3^. In the RNA silencing pathways, eAgos are loaded with short RNA fragments (guides) generated from viruses, transposons or genomic transcripts, and are directed to recognize and/or cleave target RNA for silencing. The structural basis of the RNA-guided RNA recognition of eAgos have been revealed ^4^. They consist of two lobes, which comprise N-terminal (N) and PAZ (PIWI–Argonaute–Zwille) domains, and MID (Middle) and PIWI (P-element Induced Wimpy Testis) domains, respectively ^5,6^. The MID domain contains a pocket that binds to the 5’-end of the guide RNA, while the PAZ domain anchors the 3’-end of the guide. PIWI domain forms an RNaseH fold and usually contains a DEDX (where X denotes D, H, or K) catalytic tetrad, which is essential for the RNA cleavage activity.

Prokaryotic Agos (pAgos) show much higher diversity than eAgos ^7,8^. They can be classified into three groups according to the phylogenetic analyses: long-A, long-B and short pAgos ^8^. Long-A and long-B pAgos contain all the four domains as eAgos, while short pAgos lack the N and PAZ domains. Most of long-A pAgos are active since their catalytic tetrad is intact. Many long-A pAgos are directed by DNA guides to bind and cleave DNA targets ^9–14^. In vivo, their guides are more often derived from extrachomosomal genetic elements and/or multicopy genetic elements and thus are directed to target invading viruses and plasmids ^9,14^. Long-A pAgos also acquire guides from the region of replication termination and can be directed by them to resolve replicated DNA molecules ^14,15^. Other long-A pAgos show diverse guides and targets preferences ^16–20^, suggesting that they may play versatile physiological functions.

Short pAgos are inactive as a result of the mutation of the catalytic tetrad, and their functions have been a mystery for a long time until most recently. It was reported that short pAgos and a (preduso)short pAgo from *Sulfolobus islandicus* (Si) that shares the same domain architecture as short pAgos but is not classified into the short group by the phylogenetic analysis ^21^, constitute defense systems together with their associated proteins ^22–25^. These defense systems confer immunity against viruses and plasmids via an abortive infection (Abi) response. Abi is a defense strategy that kills the infected cells or induces cell dormancy to suppress the spreading of the invaders ^26^. The general mechanism for the Abi response is that short pAgos sense invaders by guide-directed target recognition and activate their associated proteins, such as NADase and membrane protein, to induce cell death or cell dormancy ^27^.

Long-B pAgos constitute the second inactive pAgo group ^8^. A representative long-B pAgo, *Rhodobacter sphaeroides* (Rs) Ago, binds short RNA as guides and is directed to recognize target DNA ^28,29^. The DNA binding results in suppression of plasmid-encoded gene expression and/or plasmid degradation ^28,29^, but the involved mechanisms remain elusive. Moreover, gene neighborhood of long-B pAgos also encodes the so-called pAgo-associated proteins ^8^, the functional connections of which with long-B pAgos remain unknown. In this study, we explored the diversity of long-B pAgos and found that long-B pAgos are genetically and functionally associated with nucleases, NADases and membrane proteins. Relying on these associated effectors, the long-B pAgos mediate Abi response to provide immunoprotection against invading plasmid. In particular, the long-B pAgo-nuclease system performs RNA-guided unspecific DNA degradation following recognition of a specific DNA target, indicating that both CRISPR-Cas systems and pAgo systems can utilize the target recognition-activated collateral DNA degradation as a defense strategy ^30–34^.

## Results

### Long-B pAgos cluster with potential toxic effectors

Bioinformatic analysis indicates that long-B pAgos tend to associate with a number of proteins that can be clustered into several <u>o</u>rtho<u>g</u>roups (og) ^8^. Among then, og_15 (nuclease), og_44 (SIR2_2 domain containing protein) and og_100 (protein with unknown domain) are usually encoded by simple operons that only contain 2 or 3 genes ^8^. To further analyze the connection between long-B pAgos and their associated proteins, we constructed a phylogenetic tree of long-B pAgos and marked them with different colors according to their associated proteins (Figure 1A). This reveals that the long-B pAgos associated with og_15, og_44, og_54 (VirE N-terminal domain containing protein) and og_100 can be clustered into separated subclades. Further, comparison of the phylogenetic trees of og_15, og_44 and og_100, and their respective pAgos suggests the coevolution of the long-B pAgos and the associated proteins (Figure 1B-D). In addition, the operons encoding og_15, og_44 and og_100 are organized in the same structure, with *pAgos* upstream of the associated genes (Figure 1E). Together, the analyses strongly suggest that og_15, og_44 and og_100, and their respective long-B pAgos are functionally connected.

**Figure 1.**
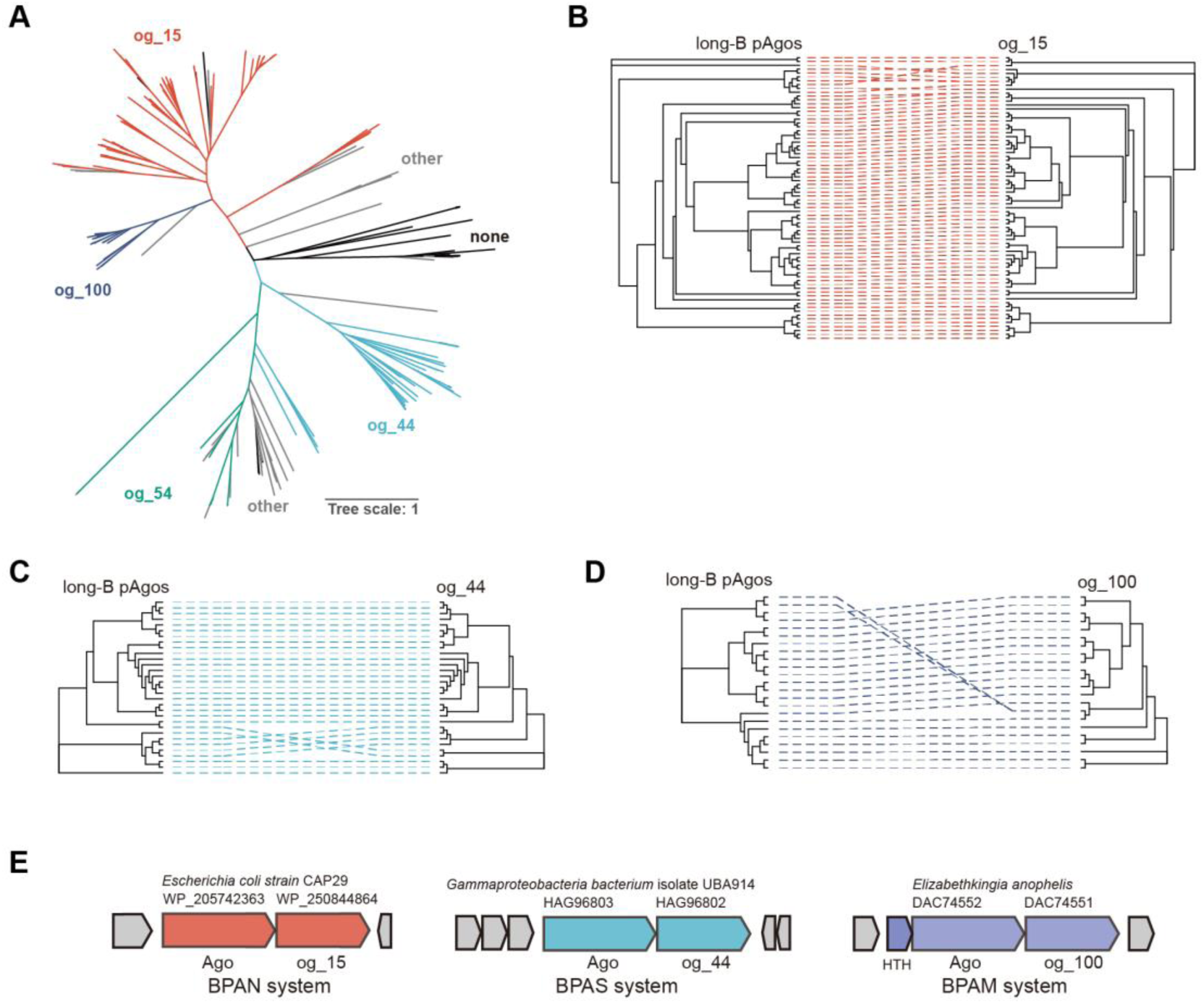
Phylogenetic analysis of long-B pAgo systems. A. Maximum likelihood-based unrooted phylogenetic tree of long-B pAgos. The proteins were derived from the supplementary data provided by Ryazansky et al., 2018, mbio. The long-B pAgos associated with different orthogroups (ogs) are marked with different color, and the corresponding ogs are indicated. none: the adjacent genes do not form an operon-like structure with the long-B *pAgo* genes; other: the adjacent genes form an operon-like structure with the long-B *pAgo* genes but do not encode any protein belonging to og_15, og_44, og_54 or og_100. B-D. Tanglegram of phylogenetic trees of og_15, og_44 and og_100 (right), and their respective long-B pAgos (left). E. Operon structure and neighborhood genes (shown in grey) of representative long-B pAgo systems. The names of the systems, their source organisms, and the encoded proteins and their accession numbers are indicated. HTH: putative helix-turn-helix transcription regulator. See also Figure S1 and Table S1.

To gain an insight into the possible function of og_100 that was annotated as “protein with unknown domain” ^8^, we predicted the structure of og_100 members using DeepTMHMM and Alphafold2. This reveals that the og_100 members are trans-membrane (TM) proteins (an example shown in Figure S1 G and H). Nucleases, SIR2-like proteins and TM proteins are widely found in Abi defense systems, including the characterized pAgo Abi systems ^22–25,35–41^. Their association with long-B pAgos implies that the related long-B pAgo systems might also mediate Abi responses. Based on the nomenclature proposed for short pAgo systems and SiAgo system ^22,25^, the long-B systems are named as BPAN (long-<u>B p</u>rokaryotic <u>a</u>rgonaute <u>n</u>uclease), BPAS (long-<u>B p</u>rokaryotic <u>a</u>rgonaute <u>S</u>ir2) and BPAM (long-<u>B p</u>rokaryotic <u>a</u>rgonaute trans-<u>m</u>embrane) systems, respectively. Correspondingly, the associated proteins are named bAgaN (long-<u>B</u> p<u>Ag</u>o-<u>a</u>ssociated <u>n</u>uclease), bAgaS (long-<u>B</u> p<u>Ag</u>o-<u>a</u>ssociated <u>S</u>ir2) and bAgaM (long-<u>B</u> p<u>Ag</u>o-<u>a</u>ssociated trans-<u>m</u>embrane). To reveal the functions and molecular mechanisms of these long-B pAgo systems, we selected the *Escherichia coli* (Ec) BPAN system, a BPAS system from unclassified *Gammaproteobacteria bacterium* (Gb) and the *Elizabethkingia anophelis* (Ea) BPAM system for analysis (Figure 1E and Table S1).

### bAgaN is a PD-(D/E)XK superfamily DNase

We began by focusing on the biochemical properties of bAgaN. bAgaN was annotated as a member of the RecB-like protein family, belonging to the PD-(D/E)XK superfamily ^8^. Members of the superfamily are involved in restriction-modification systems, DNA metabolism, tRNA splicing ^42^. Recently, many members of the superfamily are found as effectors of type III CRISPR-Cas and CBASS defense systems ^32,33^. We expressed EcbAgaN in *E. coli* BL21 and purified it to apparent homogeneity (Figure S2A). The analysis by multi-angle light scattering coupled with size exclusion chromatography (SEC-MALS) indicates that EcbAgaN forms a dimer in solution (Figure 2A). We predicted the dimer structure of EcbAgaN (Figure S1A). The analysis reveals that EcbAgaN is composed of two domains that are connected by an unstructured link and the C-terminal domain adopts a restriction endonuclease-like fold (Figure S1A and B), which possesses the conserved D-EVK motif as shown by the sequence alignment analysis (Figure S1C). A database search using the DALI structure-comparison server ^43^ reveal structural similarity to the type III CRISPR-Cas system accessory nuclease Card1 (PDB: 6wxx) ^34^ and Can2 (PDB: 7bdv) ^44^, and the type IIS restriction endonuclease R.BspD6I (PDB: 2ewf) ^45^. Then, we analyzed the nuclease activity of EcbAgaN with various substrates. The results show that EcbAgaN efficiently cleaves ssDNA, dsDNA and ssDNA from a DNA/RNA duplex but shows no activity towards RNA (Figure 2B), and the DNase activity of EcbAgaN is dependent on Mn^2+^ (Figure S2B). In addition, EcbAgaN also efficiently degrades plasmids and genomic DNA that are extracted from *E. coli* DH5α cells (Figure 2C), suggesting that EcbAgaN can cleave methylated DNA. We then constructed the mutants that carry alanine substitutions of the D-EVK motif (M1: D298A; M2: E309A-K311A). Analysis of the mutants reveals that they are inactive for DNA cleavage (Figure 2D), indicating that the motif is essential for the nuclease activity.

**Figure 2.**
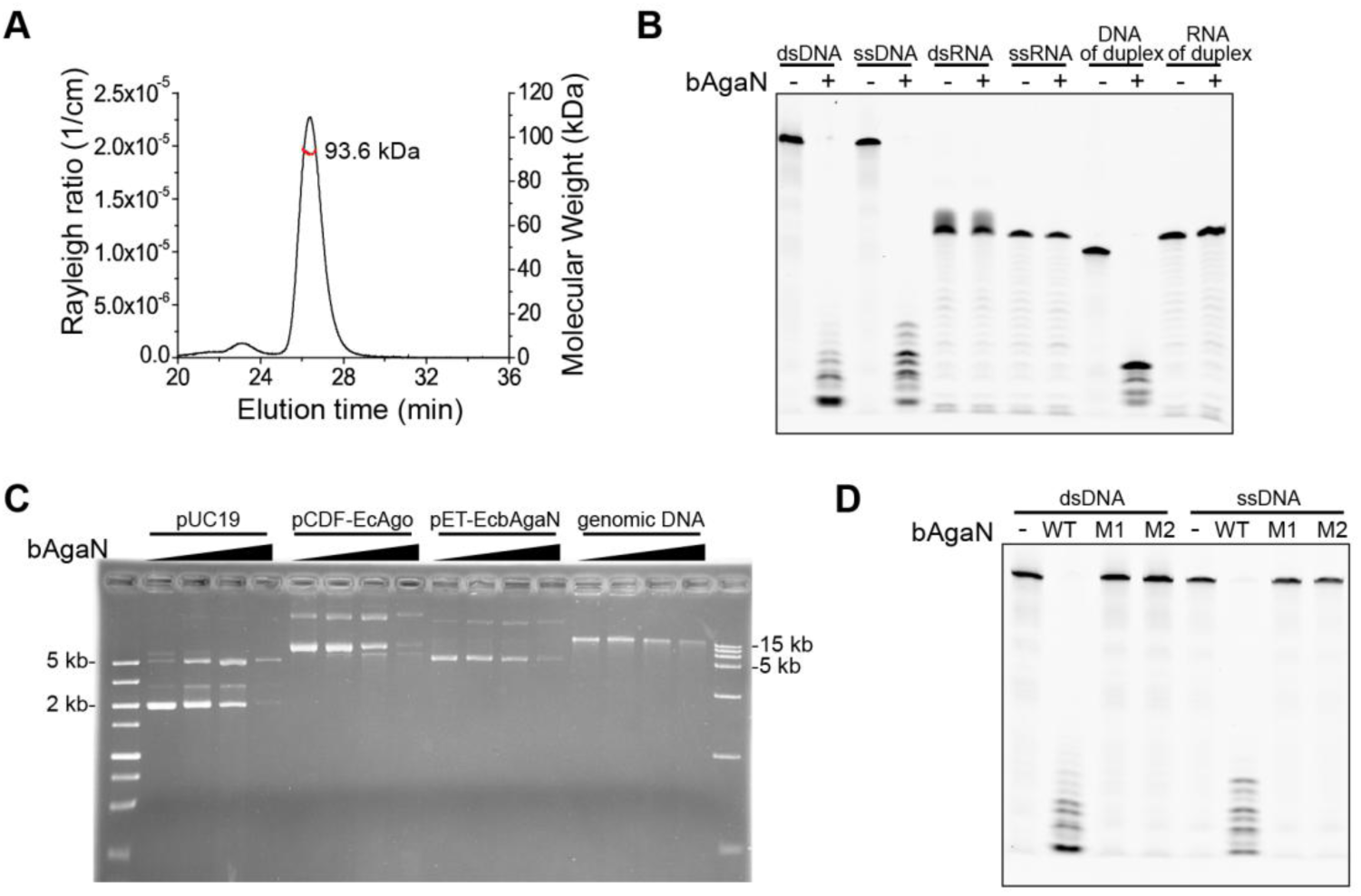
EcbAgaN is a nonspecific DNase. A. Multi-angle light scattering coupled with size exclusion chromatography (SEC-MALS) analysis of EcbAgaN. B. Substrate specificity of EcbAgaN. FAM-labeled dsDNA, ssDNA, dsRNA, ssRNA and DNA/RNA duplexes that containing either FAM-labeled DNA strand or RNA strand were incubated with EcbAgaN and then analyzed by denaturing polyacrylamide gel electrophoresis. C. Degradation of plasmid DNA and genomic DNA by EcbAgaN. Plasmid DNA, including pUC19, pCDF-EcAgo^743^, pET28T-EcbAgaN, and *E. coli* genomic DNA were incubated with a gradient of EcbAgaN and then analyzed by agarose gel electrophoresis. D. DNase activity of EcbAgaN and its mutants. M1: D298A; M2: E309A-K311A. See also Figure S1 and S2.

### The EcBPAN system confers cell toxicity via degrading genomic DNA

We noticed that EcAgo and its close homologues have two predicted starting codons, ATG and TTG, encoding a 743 amino-acids (a.a.) protein (EcAgo^743^) and a 731-a.a. Ago (EcAgo^731^) respectively (Figure S3A, Table S1). We firstly focused on the EcAgo^743^ system. We constructed strains expressing EcAgo^743^ and EcbAgaN individually or both of them (i.e., the BPAN system) using the expression plasmids pCDF-EcAgo^743^ and pET28T-EcbAgaN (Table S2) respectively or together. The strains, as well as the control strain containing empty vectors (EV), were grown in the presence of inducers (IPTG and aTc) but without antibiotics. At 4 h during the induction, cell viability was assessed by plating of the cells onto plates with or without antibiotics. The results derived from the antibiotic-free plates show that individual expression of EcAgo^743^ and EcbAgaN has little effect on the cell viability, while expression of the EcBPAN system results in an over 100-fold reduction in the cell viability (Figure 3A). The results indicate that EcAgo^743^ and EcbAgaN cooperatively confer cytotoxicity. In addition, the mutated EcBPAN systems containing EcbAgaN dead mutants (M1 and M2) do not induce any reduction in the cell viability, indicating that the cytotoxicity is dependent on the nuclease activity of EcbAgaN.

**Figure 3.**
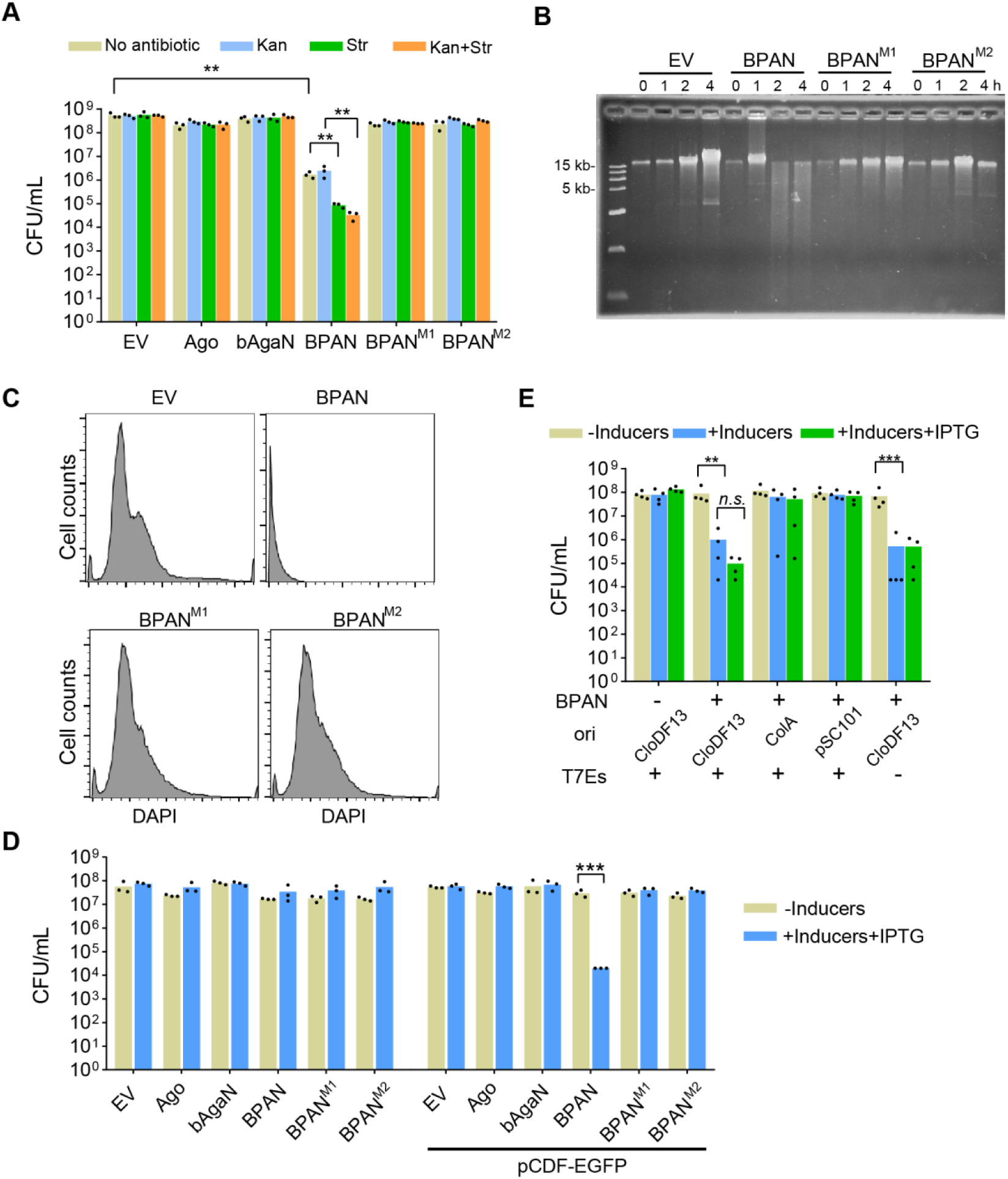
EcBPAN system is activated by the CloDF13 origin and degrades genomic DNA. A. Expression of the EcBPAN system triggers cell death. The cells containing empty vector (EV), EcAgo, EcbAgaN, the EcBPAN system and the mutated EcBPAN systems (M1 and M2) were grown in LB medium supplemented with IPTG and aTc for 4 h. Then, the cells were plated onto the plates with or without antibiotics as indicated. The colony formation unit per mL (CFU/mL) was calculated. B. EcBPAN system degrades genomic DNA in vivo. After the wild type and mutated EcBPAN systems were induced by IPTG and aTc, genomic DNA was extracted at indicated time points and analyzed by agarose gel electrophoresis. C. Flow cytomerty analysis of DNA content distributions in the cells after the wild type and mutated EcBPAN systems were induced by IPTG and aTc for 2h. D. The pBAD-expressed EcBPAN system is activated by pCDF-EGFP. The cells expressing EcAgo, EcbAgaN, the EcBPAN system and the mutated EcBPAN systems (M1 and M2) in the presence or absence of pCDF-EGFP were plated onto the plates without inducers or with the inducers plus IPTG. Inducers: arabinose and aTc. E. The pBAD-expressed EcBPAN system is activated by the CloDF13 origin. The cells carrying EcBPAN system were transformed with pCDF-EGFP and its variants, where the CloDF13 origin was substituted with the indicated origins or the T7 expression cassettes (T7Es) were removed. Then, the cells were plated onto the plates with or without inducers, or with inducers and IPTG. Inducers: arabinose and aTc. For all bar graphs, CFU/mL was calculated and the average of three or four biological replicates are shown, with individual data points overlaid. **: p < 0.01; ***: p < 0.001; n.s.: not significant. See also Figure S3.

To reveal how the EcBPAN system mediates cytotoxicity, we analyzed the genome integrity of the cells. The cell samples were taken from the cultures during the induction and subjected to genomic DNA extraction and DAPI-staining. Gel electrophoresis analysis of the genomic DNA indicates that the EcBPAN system results in extensive genomic DNA degradation at 2 h (Figure 3B). Meanwhile, analysis of the DAPI-stained cells with flow cytometry shows that the EcBPAN system induces dramatic decrease of the cellular DNA content (Figure 3C). By comparison, the dead mutations of EcbAgaN abolished the observed genomic DNA degradation (Figure 3B and C). Together, the data indicate that EcAgo^743^ and EcbAgaN cooperatively induce cytotoxicity by degrading genomic DNA with the nuclease activity of EcbAgaN.

### The cell toxicity of the EcBPAN system is triggered by the CloDF13 origin

To reveal whether the system could also induce plasmid depletion, we compared the cell viability on the antibiotic-free plates and the plates containing Kanamycin (Kan, for selection of pET28aT-EcbAgaN) or streptomycin (Str, for selection of pCDF-EcAgo^743^), or both of them (Figure 3A). The results show that individual EcAgo^743^ or EcbAgaN, or the mutated EcBPAN systems do not lead to any plasmid depletion. By comparison, the wild type EcBPAN system induces a 100-fold decrease in the cell viability on the Str plates compared to the antibiotic-free plates, and on the Kan+Str plates compared to the Kan plates (Figure 3A), indicative of the depletion of the pCDF-EcAgo^743^ plasmid. Nevertheless, the cell viability on the Kan plates is the same as that on the antibiotic-free plates. The results suggest that the EcBPAN system might mediate selected depletion of pCDF-EcAgo^743^. To explore the phenomenon, we expressed EcAgo^743^ and EcbAgaN using the pBAD24 plasmid with araBAD promoter and Tet promoter respectively. In this case, induction of EcAgo^743^ and EcbAgaN using arabinose and aTc does not induce cell viability reduction (Figure 3D). Then, the cells were transformed with the pCDF-EGFP plasmid, which, however, rendered the expression of EcAgo^743^ and EcbAgaN toxic, resulting a ∼1000-fold reduction in the cell viability (Figure 3D). In addition, the cell viability reduction requires both EcAgo^743^ and the nuclease activity of EcbAgaN. The data indicate that pCDF-EGFP activates the EcBPAN system to mediate cell death. Thus, the depletion of pCDF-EcAgo^743^ in Figure 3A could be due to the selective killing of the cells carrying the plasmid.

The CloDF13 origin and T7 expression cassettes of pCDF-EGFP have been shown to act as triggers for short pAgo systems ^23,25^. To gain further insights into how pCDF-EGFP triggers the toxicity of the EcBPAN system, we replaced the CloDF13 origin of pCDF-EGFP with the ColA origin and the pSC101 replicon respectively, and also removed the T7 expression cassettes (T7Es) of pCDF-EGFP. Then, the cell viability of the cells containing these variant plasmids was analyzed on the plates in the presence or absence of IPTG (Figure 3E). The results indicate that the origin replacement abolishes the cytotoxicity of the EcBPAN system, while removal of T7Es has little effects on the cytotoxicity. In addition, IPTG has little effects on the cytotoxicity, either. Together, the data indicate that the CloDF13 origin activates the EcBPAN system to trigger cell death. We also analyzed the functions of the EcBPAN system expressing EcAgo^731^. The results show that the EcAgo^731^ system does not induce any reduction in the cell viability when expressed from the pBAD24 vector, while pCDF-EGFP activates the cytotoxicity of the system (Figure S3B), indicating that presence of the 12 a.a. or not does not alter the functions of EcAgo.

Next, we analyzed whether the EcBPAN system could provide immunoprotection against phages. Efficiency of plaque formation (EOP) of T5, T7 and lambda-vir on the bacterial lawn expressing the EcBPAN system or its mutants was measured (Figure S3C). The results show that the EcBPAN system did not induce any reduction in the EOP of any phage.

### In vivo nucleic acid binding of EcBPAN system

pAgo proteins are inherently directed by nucleic acid guides to recognize complementary nucleic acid targets ^9,28^, and the ability is employed by short pAgo and SiAgo systems to sense invaders ^22,23,25^. To gain insight into how the EcBPAN system sense the invading plasmid, we analyzed the nucleic acids associated with EcAgo in vivo. Specifically, EcAgo^743^ was purified from the cells containing pCDF-EGFP (+pCDF) or EcbAgaN (+bAgaN) or both of them (+pCDF, +bAgaN), and the nucleic acids were extracted from the three protein samples. The nucleic acids were subjected to Fast Thermosensitive Alkaline Phosphatase (FastAP) treatment or not, followed by T4 polynucleotide kinase (PNK) labeling with γP^32^-ATP. Only the FastAP-treated nucleic acids were efficiently labeled by PNK (Figure 4A), indicating that most of the nucleic acids contain 5’ phosphate group. The assay also indicates that the nucleic acids were less abundant in the sample lacking pCDF-EGFP. Then, treatment of the labeled nucleic acids by RNase and DNase respectively reveals that only RNase degraded the nucleic acids (Figure 4B). Together, the EcAgo-associated nucleic acids are small RNAs containing 5’ phosphate group.

**Figure 4.**
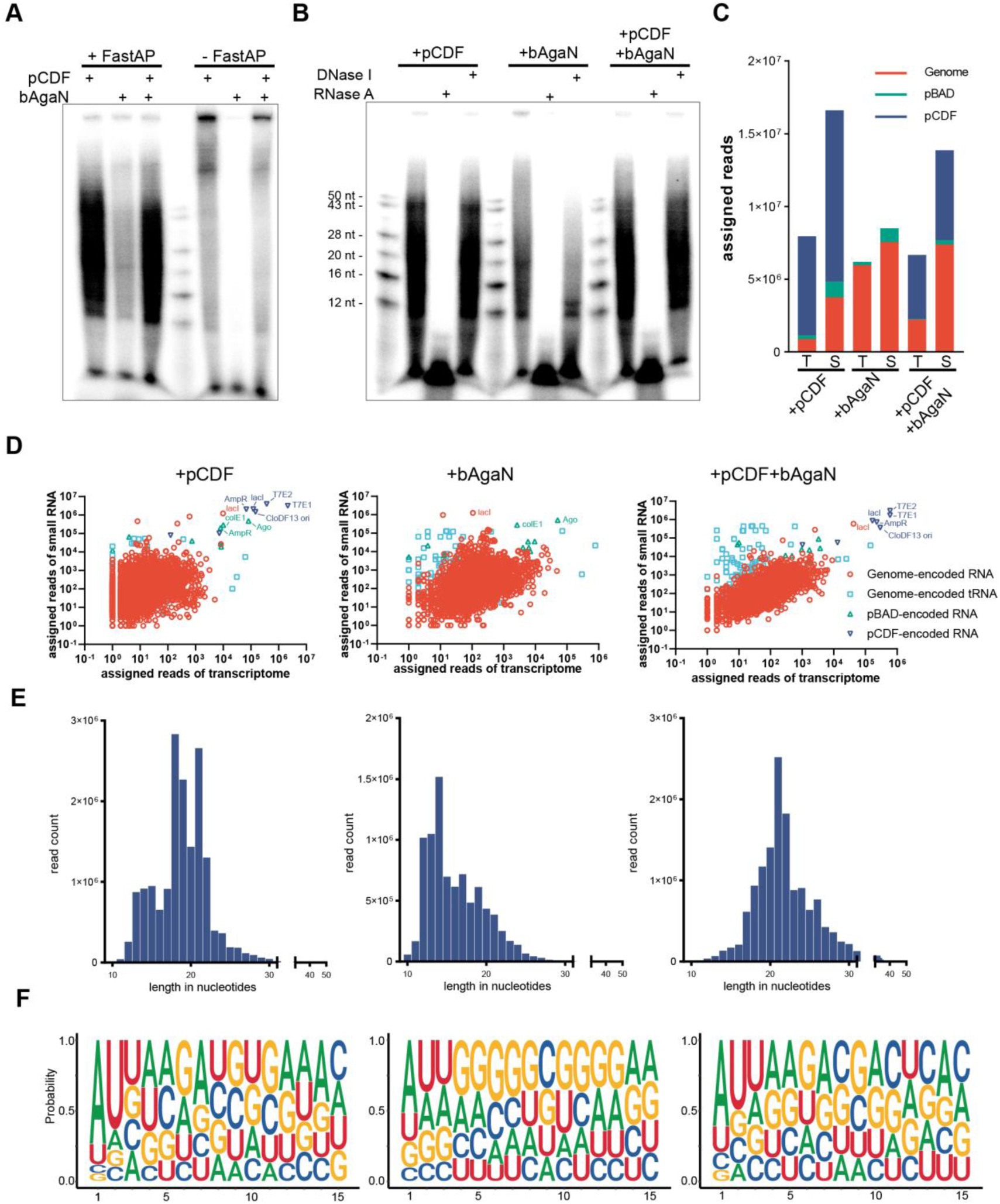
EcAgo associates with small RNAs that are derived from transcriptome and contain 5’-end phosphate group. A. Analysis of the 5’-end of EcAgo-associated nucleic acids. Nucleic acids were extracted from the EcAgo^743^ protein samples that were purified from the cells containing pCDF-EGFP or EcbAgaN or both. The nucleic acids were treated with FastAP or not, and then labeled with γP^32^-ATP, followed by analysis by denaturing polyacrylamide gel electrophoresis. B. Treatment of the labeled nucleic acids with DNase I and RNase A, respectively. C. Reads numbers of the transcriptome sequences (T) and small RNA sequences (S) that are assigned to genome and plasmids. pBAD: pBad24-EcAgo^743^ or pBad24-EcAgo^743^–EcbAgaN; pCDF: pCDF-EGFP. D. Correlation between the transcriptome sequences and small RNA sequences. The Pearson correlation coefficients of the three samples are r > 0.65, r > 0.034, r > 0.88, respectively, with p values < 10^−99^, < 0.03, < 10^−99^, respectively. E. Length distribution of the small RNAs. F. Nucleotide bias of the small RNAs. See also Figure S4 and S5, and Table S5.

Next, the small RNAs were subjected to RNA sequencing, and meanwhile the transcriptome of the corresponding cultures was also analyzed. The results show that the abundance of small RNAs is correlated with the RNA sequences in the transcriptome in the samples containing pCDF-EGFP (Figure 4C and D, Table S5). Moreover, pCDF-EGFP generated much more RNAs than the pBAD24 plasmids and even more than the *E. coli* genome in the transcriptome. Consequently, the pCDF-EGFP-derived small RNAs are much more abundant that those derived from the pBAD24 plasmids and the genome (Figure 4C and D), in agreement with that only pCDF-EGFP activates the EcBPAN system. On the other hand, absence of EcbAgaN has no apparent effects on the abundance of the small RNAs or their correlation with the transcriptome.

Then, we analyzed the length distribution and sequence bias properties of the small RNAs. From the cells containing pCDF-EGFP, the small RNAs are predominantly ∼18-22 nt in length and show a bias toward A and U at the first two nt, independent of EcbAgaN (Figure 4E and F). By comparison, absence of pCDF-EGFP lowered the AU bias and results in the accumulation of the small RNAs shorter than 18 nt, possibly because the genome-derived small RNAs are shorter than the plasmid-derived small RNAs (Figure S4). Intriguingly, the length distribution and sequence bias properties are applied to the small RNAs derived from both the CloDF13 origin and other genetic elements that do not activate the EcBPAN system, such as the pBAD24 plasmids and the two T7 expression cassettes (T7Es) of pCDF-EGFP (Figure S4 and S5). In particular, T7Es generate even more abundant small RNAs than the CloDF13 origin. The data suggest that EcAgo does not specifically acquire small RNAs from the CloDF13 origin-derived transcripts, although it is the specific trigger.

### Guide-directed target recognition of EcAgo activates EcbAgaN

To gain further insight into how the EcBPAN system senses invading genetic element and how it is activated, we performed an electrophoretic mobility shift assay (EMSA) to analyze the guide loading and target binding properties of EcAgo. Both EcAgo^743^ and EcAgo^731^ were successfully purified (Figure S2C). However, binding by EcAgo^743^ resulted in that the nucleic acid substrates were stuck in the wells of the gel (data not shown). Thus, we analyzed the nucleic acid binding properties of EcAgo^731^. Single strand (ss) RNA and DNA substrates containing a 5’ phosphate (5P) group and a 5’ hydroxyl group (5OH) respectively were used in the assays. The results show that EcAgo^731^ efficiently binds to 5P-RNA (Figure 5A), in agreement with that it associates with 5P-RNA in vivo. Moreover, when preloaded with 5P-RNA as a guide, EcAgo^731^ can specifically recognize target ssDNA, rather than non-target ssDNA, target RNA or double strand DNA containing the target sequence (Figure 5B). This suggests that although not detected in the EcAgo-copurified nucleic acids, ssDNA could serve the target of the EcBPAN system. We further analyzed whether EcAgo could mediate any DNA cleavage. Incubation of the ssDNA and dsDNA substrates with apo EcAgo^731^ or the EcAgo^731^ supplemented with guide RNA and/or target DNA does not result in any cleavage of the substrates (Figure S2D), suggesting that the mutation of the catalytic site indeed inactivates the long-B pAgo.

**Figure 5.**
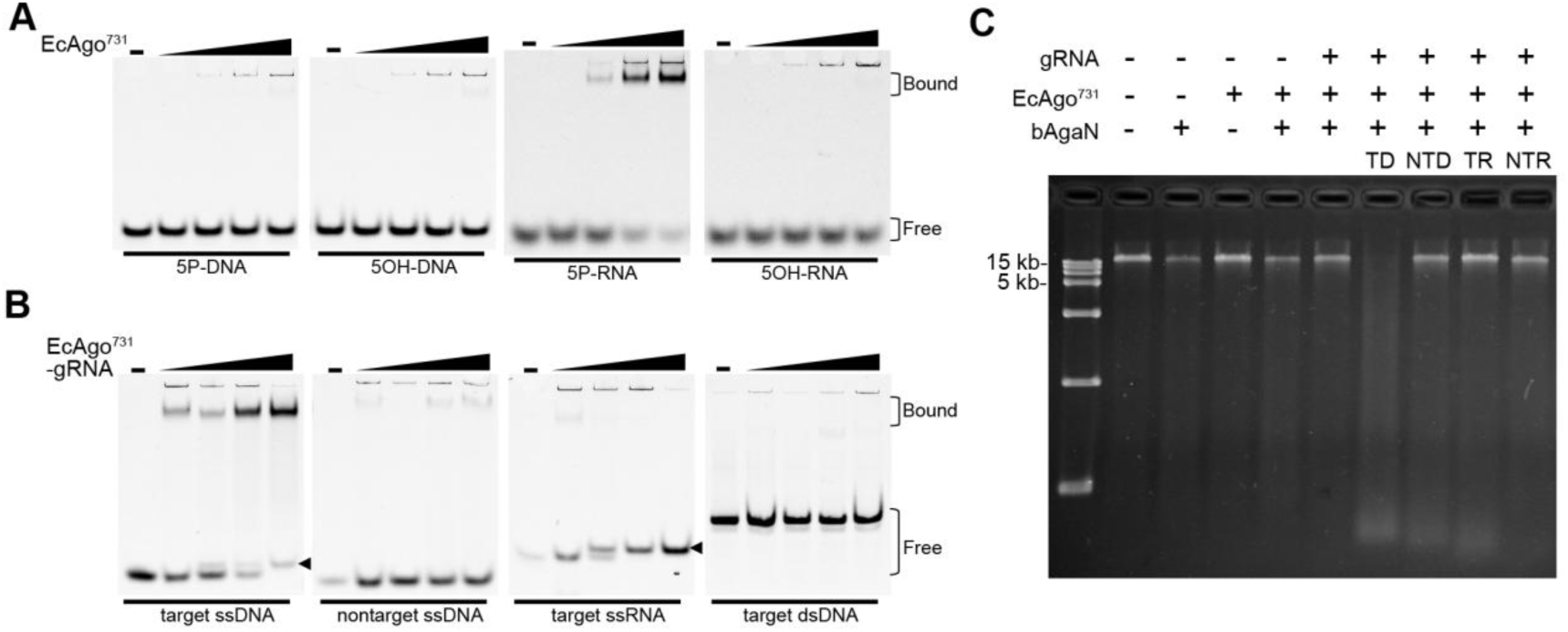
EcAgo is directed by 5P-RNA guides to bind ssDNA targets and activates EcbAgaN. A. Nucleic-acid binding of EcAgo^731^. The different substrates are indicated below the panels. B. Target-binding of EcAgo^731^ preloaded with 5P-RNA as guide (gRNA). The different substrates are indicated below the panels. The arrows indicate duplex formed by the guide and target ssDNA or ssRNA. C. Target recognition of the gRNA-bound EcAgo activates EcbAgaN. About 200 ng genomic DNA was treated with EcbAgaN. Aliquots of the reaction were also supplemented with EcAgo^731^, 5P-RNA and/or target ssDNA or other oligonucleotides. TD: target ssDNA; NTD: nontarget ssDNA; TR: target ssRNA; NTR: nontarget ssRNA. See also Figure S2 and S6.

The above data suggest that RNA-directed target DNA binding by EcAgo may activate the system. To constitute the activation in vitro, the EcAgo^731^–gRNA (5P-RNA) complex was incubated with target ssDNA, nontarget ssDNA, target RNA or nontarget RNA. The resulting mixtures, as well as apo EcAgo^731^ and the EcAgo^731^–gRNA complex, were analyzed for their effects on the DNase activity of EcbAgaN using genomic DNA, plasmid DNA and PCR product as substrates (Figure 5C and Figure S6). The results show that the DNase activity of EcbAgaN is significantly stimulated by the EcAgo^731^–gRNA complex supplemented with the target ssDNA instead of other oligonucleotides. The data reveal that target ssDNA recognition of guide-directed EcAgo activates EcbAgaN for nonspecific DNA degradation.

### The GbBPAS system is activated by the CloDF13 origin to mediate NAD^+^ depletion

Next, we aimed to reveal whether BPAS systems also provide immunity against invading plasmid. bAgaS has been annotated as a Sir2_2-domain-containing protein and is only distantly related to the Sir2 domain of the short pAgo-associated Sir2-APAZ protein ^8^. Structural prediction reveals that the Sir2_2 of domain GbbAgaS is similar to that of ThsA protein ^46^ and they share the conserved residues of the NAD-binding site (Figure S1D-F). We constructed strains expressing individual GbAgo and GbAgaS proteins, GbBPAS system as well as the GbAgaS mutant (N155A) protein or system using the pBAD24 vector. Alanine substitution of the Asparagine in ThsA can abolish its activity and function ^46,47^. To analyze the potential function of pCDF-EGFP, the above-mentioned strains were also transformed with the plasmid. Then, the strains with or without pCDF-EGFP were plated onto the plates containing corresponding antibiotics and with or without the inducer (arabinose), and the cell viability was measured (Figure 6A). The results show that individual GbAgaS induces a ∼10^4^-reduction in the cell viability, which is independent of pCDF-EGFP but is abolished by GbAgo. On the other hand, GbAgaS and GbAgo only confer cytotoxicity in the presence of pCDF-EGFP, indicating that the plasmid activates the GbBPAS system. In addition, the N155A mutation abolishes the reduction of cell viability, indicating that the activity of the Sir2_2 domain is essential for the cytotoxicity. The variants of pCDF-EGFP were also tested for their ability to activate the GbBPAS system (Figure 6B). This reveals that the CloDF13 origin activates the GbBPAS system as observed for the EcBPAN system.

**Figure 6.**
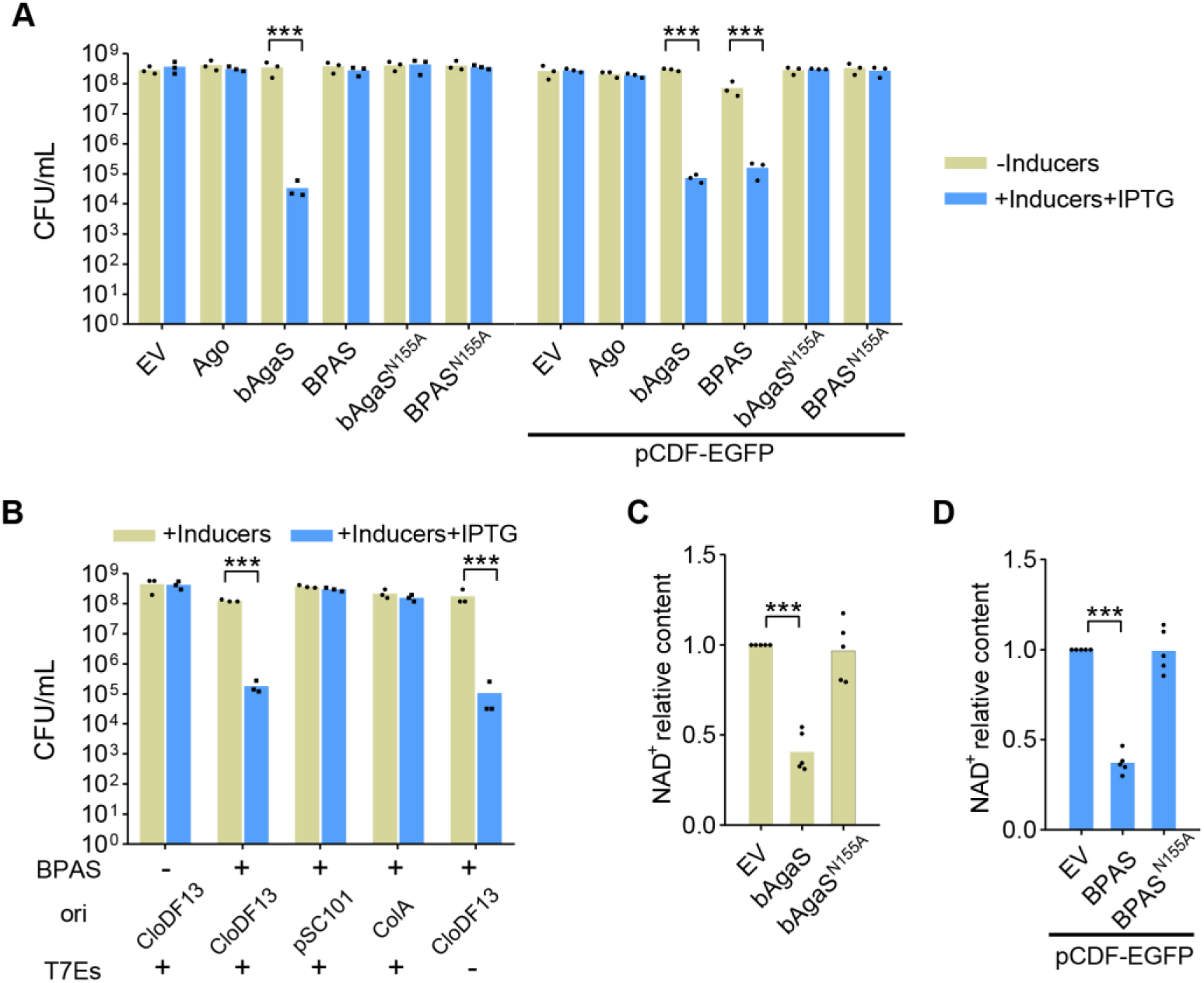
GbBPAS system is activated by the CloDF13 origin and mediates NAD^+^ depletion. A. GbBPAS system is activated by pCDF-EGFP to induce cell death. The cells containing empty vector (EV), the wild type and mutant GbBPAS systems, and individual proteins as well as mutants in the presence or absence of pCDF-EGFP were plated onto the plates without inducer or with the inducer and IPTG. Then, CFU/mL was calculated. B. Activation of the GbBPAS system requires the CloDF13 origin. Cell viability of the cells carrying the GbBPAS system in the presence of pCDF-EGFP or its variants was measured. C. GbbAgaS reduces cellular NAD^+^ level. NAD^+^ amount of the cells carrying wild type and mutant GbbAgaS proteins was measured, and relative NAD^+^ level was calculated with the values of the EV samples set as 1. D. GbBPAS system reduces cellular NAD^+^ level in the presence of pCDF-EGFP. NAD^+^ amount of the cells carrying wild type and mutant GbBPAS systems was measured, and relative NAD^+^ level was calculated. For all bar graphs, the average of three or five biological replicates are shown, with individual data points overlaid. ***: p < 0.001; inducer: arabinose. See also Figure S7.

Since many defense systems containing Sir2-like domains confer immunity by NAD^+^ depletion ^23,24,46,47^, we analyzed whether the GbBPAS system can also deplete NAD^+^. The cells expressing GbAgaS or its mutant, the GbBPAS system or the mutated system in the presence of pCDF-EGFP, were grown in liquid medium and protein expression was induced by arabinose. The results show that GbAgaS individually and the GbBPAS system together with pCDF-EGFP induce significant culture growth retardation and NAD^+^ level reduction (Figure S7A and B, Figure 6C and D), in line with the cell viability results. Moreover, the N155A mutation abolishes the reduction in NAD^+^ level and relieves the growth retardation. Together, The data indicate that the GbBPAS system confers immunity against invading plasmid via the Sir2_2 domain-mediated NAD^+^ depletion.

### The EaBPAM system triggers cell death depending on the trans-membrane (TM) effector

The functions of the EaBPAM system were analyzed using the same methods as applied for the GbBPAS system. Instead of a site mutation, we constructed a truncation mutant of EabAgaM (EabAgaM^ΔTM^) that lacks the C-terminal TM region (106 a.a.) (Figure S1G and H). The results show that the EaBPAM system triggers a ∼10^3^-reduction in the cell viability, which requires both EaAgo and the full-length EabAgaM (Figure 7A). Nevertheless, the pCDF-EGFP plasmid is not required since the system expressed from the pBAD24 vector confers cytotoxicity in the absence of pCDF-EGFP. Then, we analyzed the cytotoxicity of the system expressed from pKD46, a low-copy vector. Again, the EaBPAM system induces significant cell death regardless of the presence of other plasmids (Figure 7B). When expressed in liquid medium, the EaBPAM system results in growth retardation (Figure S7C). However, it does not induce significant membrane depolarization or membrane destruction (Figure S7D and E) as other defense systems carrying a TM effector usually do ^22,26,37^.

**Figure 7.**
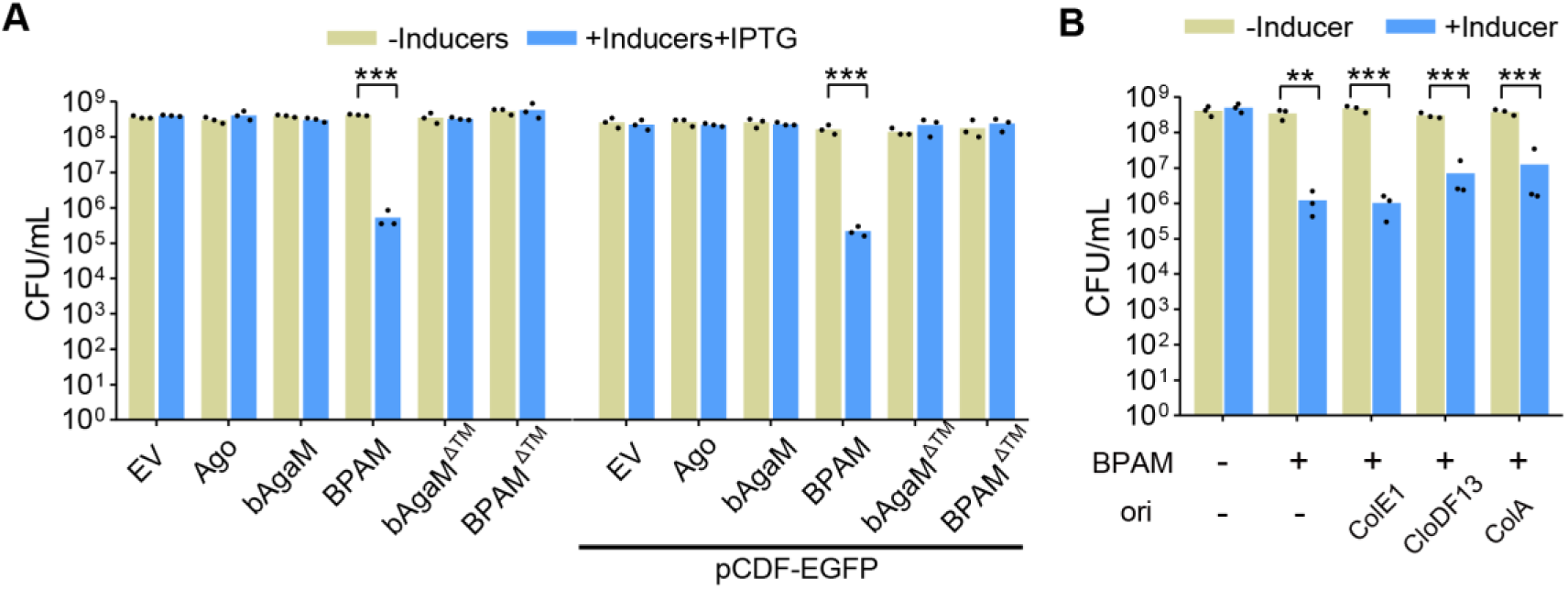
EaBPAM system induces cell death. A. Cell viability of the cells containing empty vector (EV), the wild type and mutant EaBPAM systems, and individual proteins as well as mutants in the presence or absence of pCDF-EGFP was analyzed. B. Cell viability of the cells containing the EaBPAM system in the presence of pCDF-EGFP or its variants was measured. **: p < 0.01; ***: p < 0.001; inducer: arabinose. See also Figure S7.

## Discussion

In this study, we characterized three different long-B pAgo systems that comprise ∼60% of all long-B pAgos (Figure 1A). The three systems are equipped with different associated effectors respectively, and can employ the effectors to induce cell death. Two of them, i.e. the BPAN and BPAS systems, are activated by the CloDF13 origin to mediate genome degradation and NAD^+^ depletion, respectively. The genome degradation is performed by the nuclease of the BPAN system, which is activated by the target DNA recognition of the RNA-guided long-B pAgo. With such a mechanism, the systems can defend against invaders via Abi responses.

Abi is the second most prevalent defense strategy that is manifested by many defense systems ^27,48^. Abi provides population-level immunity by killing the infected cells or triggering cell dormancy, which can suppress the spreading of the invader, gain time for other immune systems to clear the invader, and finally remove the invader from the cell population ^26,27^. Indeed, the EcBPAN system can specifically deplete the invader plasmid (pCDF), while the innocent plasmid (pET28T) is retained (Figure 3A). The phenomena could be due to the selective killing of the cells containing the invader after protein expression was induced, while the cells depleting the invader can survive.

Any defense system has to specifically discriminate invaders from self, while the guide-directed target recognition ability of pAgos is supposed to achieve such a function ^22,23,25^. Remarkably, all characterized inactive pAgos, including EcAgo and RsAgo as representative long-B pAgos and several representative short pAgos, prefer small RNAs containing a 5’-end phosphate group (5P-RNA) as guides and single strand DNA as target (Figure 5) ^23,25,28,29^. Moreover, they associate with the 5P-RNA guides in vivo and the abundance of the guides are correlated with that of the RNAs in transcriptome (Figure 4) ^25,28^. This suggests that the high abundance of the RNAs derived from invaders may contribute to the invader discrimination.

Nevertheless, the abundance of the genome-derived guides is comparable to that from the invaders ^25,28^ (Figure 4), indicating that the invader discrimination does not only occur at the guide-acquisition step. EcBPAN, GbBPAS system and *Geobacter sulfurreducens* short pAgo system (GsSPARSA) are specifically activated by the CloDF13 origin, instead of other genetic elements from the same plasmid (Figure 3 and 6)^23^, even though the latter, e.g. the T7 expression cassettes, may generate more guides than the CloDF13 origin as observed for EcAgo. CloDF13 origin, belonging to the ColE1-like type, uses two RNAs to initiate plasmid replication and generates ssDNA region ^49,50^, which possibly provides available target DNA. Indeed, for *Maribacter polysiphoniae* short pAgo system (MapSPARTA), the high copy number of invading plasmid plays a role in the activation of the Abi response possibly by increasing the accessibility of the target DNA ^25^. On the other hand, the ColE1 origin that initiates plasmid replication with a similar mechanism does not activate EcBPAN system or GbBPAS system. The reason for the phenomena might be that the ColE1 origin generates much less guides than the CloDF13 origin (Figure 4D). In contrast, the EaBPAM system, even when expressed from a low-copy plasmid pKD46 with a different replication mechanism ^51^, still induces cell death. Together, these studies demonstrate the diversity of the invader discrimination mechanisms of these inactive pAgo systems.

The three long-B pAgo systems employ different effectors to mediate cytotoxicity, i.e. the nuclease EcbAgaN, the Sir2_2-domain-containing protein GbbAgaS and the trans-membrane (TM) protein EabAgaM, respectively. Interestingly, these effectors are commonly found in many Abi systems ^22–25,35–38^, suggesting that the defense systems can exchange their effectors during the evolution history. EcbAgaN is activated by the specific target recognition of guide-directed EcAgo and performs indiscriminate DNA degradation to mediate Abi. The immune process resembles the collateral DNA degradation of some CRISPR-Cas systems after crRNA-directed target binding ^31–34^, indicative of the convergent evolution of the two nucleic acid-directed defense systems. Similar to the CRISPR-Cas systems, the EcBPAN system also has potential to be repurposed for nucleic acid detection using DNA substrates ^30^ or other sequence-specific technologies.

The GbBPAS system mediates NAD^+^ depletion in the presence of the invading plasmid via the Sir2_2 domain of GbbAgaS, in agreement with the canonical functions of Sir2-like domains in many defense systems ^23–25,47^. Based on our findings of the EcBPAN system and previous studies of the SPARSA systems ^23^, it is tempting to suggest that the GbBPAS system is also activated by the target recognition of the guide-bound GbAgo. Interestingly, GbbAgaS alone can induce significant cytotoxicity and the cytotoxicity is suppressed by GbAgo. The regulation is in a similar manner to the TIR-APAZ protein of SPARTA systems ^25^. By comparison, EcbAgaN and EabAgaM do not reduce cell viability individually, indicative of diverse regulation mechanisms of the associated effectors. In addition, although the EaBPAM system efficiently induces cell death, the system does not mediate membrane depolarization or membrane destruction as other TM effectors usually do ^22,26,37^. The specific mechanism how EabAgaM mediates cytotoxicity remains to be analyzed in future.

In conclusion, we demonstrate that long-B pAgo systems generally confer immunity via Abi using their associated proteins as toxic effectors. During the immune response, pAgo senses invaders by RNA-directed target DNA recognition and activates the associated effector. On the other hand, our phylogenetic analysis reveals that a minority of long-B pAgos cluster with other proteins, such as the og_54 VirE-N domain-containing protein, or do not have any associated protein in an operon (Figure 1A). The biological functions and mechanisms of these long-B pAgos need to be addressed by future studies.

## Acknowledgments

The research was supported by National Key Research and Development program of China (2022YFA0912200, 2019YFA0906400), National Natural Science Foundation of China (Grant No. 31970545, 32270099, and 31970544), Fundamental Research Funds for Central Universities (Grant No. 2662020SKPY001), the Foundation of Hubei Hongshan Laboratory (No. 2021hszd022) and Huazhong Agricultural University Scientific & Technological Self-innovation Foundation. We thank the core facilities of Center for Protein Research (CPR) and Experimental Teaching Center of Bioengineering at Huazhong Agricultural University for technical support.

## Author contributions

X.S., S.L. (Sheng Lei), S.H.L. (Shunhang Liu) and Y.L. conducted the experiments with the assistance from P.F., Z.Z., K.Y. and Y.C.. Y.L. performed phylogenetic analysis and RNA sequencing analysis. M.L. and Q.S. gave important advice and critically commented the draft. W.H. acquired the funding, supervised the work and wrote the original draft. All authors contributed to review and editing.

## STAR METHODS

### RESOURCE AVAILABILITY

#### Lead Contact

Further information and requests for resources and reagents should be directed to and will be fulfilled by the Lead Contact, Wenyuan Han.

#### Materials Availability

Plasmids, strains and other unique reagents generated in this study are available upon request.

#### Data and Code Availability

- All raw data from the assays reported in this paper will be shared by the lead contact upon request. In addition, small and total RNA sequencing data are available on the NCBI Sequence Read Archive under BioProject ID PRJNA932940.
- This study did not generate any unique code.
- Any additional information required to reanalyze the data reported in this work paper is available from the lead contact upon request.

### EXPERIMENTAL MODELS AND SUBJECT DETAILS

*Escherichia coli* DH5a were routinely grown in Lysogeny broth (LB) medium and used for plasmid cloning, while *E. coli* BL21 (DE3) was used for protein expression and the in vivo assays. *E. coli* phages T5, T7 and Lambda-vir are gifts from Shi Chen lab (Wuhan University, China).

### METHOD DETAILS

#### Plasmid construction

In general, the plasmids were constructed with the restriction-ligation cloning method or assembled with ClonExpress II One Step Cloning Kit (Vazyme, Nanjing, China). For the former, the gene fragments were amplified using the primers containing restriction sites (Table S3), digested by restriction enzymes and inserted into the plasmid vectors between the corresponding sites. For the latter, the fragments of gene coding sequences, replication origins, and expression cassettes were amplified with the primers that have homologous sequences with the backbone vectors, and the linear vectors were prepared either by restriction digestion or by PCR. Then, the fragments were assembled following the instruction of the manufacturer. The plasmids listed in Table S2 were used as vectors to express long-B pAgos and their associated proteins and to construct the trigger plasmids. The primers used for plasmid construction were listed in Table S3 with the cloning methods indicated. All cloning work was performed with *E. coli* DH5α. All the primers were synthesized by Tingke (Beijing, China).

Specifically, the optimized coding sequences of long-B pAgos and their associated proteins were synthesized by Tingke (Beijing, China). In particular, the TTG start codon of EcAgo^731^ was replaced with ATG. To obtain EcAgo and EcbAgaN proteins for biochemical analysis, the coding sequences of EcAgo^743^ and EcAgo^731^ were inserted into pCDFDuet-1, while the coding sequence of EcbAgaN were inserted into pET28a, allowing the proteins were expressed from an IPTG-induced promoter with N-terminal 6×His tag. Site mutagenesis was performed by overlapping PCR.

For in vivo experiments, pET28T, where the T7 promoter and lacO operator were replaced by a TetR expression cassette and a Tet promoter from the p46Cpf1-OP2 plasmid (Addgene #98592), was used to express EcbAgaN and its mutants, as well as His-tag-free EcbAgaN (pET28T-EcbAgaN_HF). In addition, the coding sequences of EcAgo^743^ and EcAgo^731^ were also inserted into pBAD24 to express the proteins under control of an araBAD promoter. To express EcBPAN system using pBAD24, a fragment containing TetR expression cassette, Tet promoter, EcbAgaN coding sequence and the terminator was amplified and inserted into pBAD-EcAgo^743^. The plasmids expressing mutant EcBPAN systems were constructed in a similar way.

To generate the pCDF-ΔT7 plasmid, the DNA fragment lacking T7 expression boxes was amplified from pCDF-EGFP. Then, recircularization of the fragment was performed by ClonExpress II One Step Cloning Kit. To replace the origin of pCDF-EGFP plasmids with ColA or pSC101^TR^, the DNA fragments containing ColA or pSC101^TR^ replicons, and the backbone of pCDF-EGFP plasmid were amplified respectively, and assembled using the ClonExpress kit.

For the in vivo experiments of GbBPAS system and EaBPAM system, the coding sequences of long-B pAgos and their associated proteins were inserted into pBAD24 respectively. Then, the expression cassettes of GbbAgaS and EabAgaM were amplified and inserted into pBAD24-GbAgo and pBAD24-EaAgo, respectively. The site mutagenesis of GbbAgaS was performed by overlapping PCR, while the truncation of *EabAgaM* gene was performed by amplification of the fragment lacking the gene C-terminal region from pBAD24-EaAgo-EabAgaM and pBAD24-EabAgaM respectively and recircularization of the fragment.

To construct the pKD46-M1-EaAgo-EabAgaM plasmid, the p^TR^Cas12a-NT plasmid from our previous study ^52^ was used as template to amplify the Chl-resistant gene and the araBAD promoter-araC-pSC101 origin fragment. Then, the fragments were assembled with the fragment containing *EaAgo* and *EabAgaM* genes amplified from pBAD24-EaAgo-EabAgaM.

#### Protein expression and purification

The plasmids used for purification of EcAgo and EcbAgaN proteins and their variants were transformed into *E. coli* BL21(DE3). The transformants were grown in LB medium at 37 °C containing the corresponding antibiotics. At an optical density (OD_600_) of ∼1.0, the cultures were cooled on ice for 10 min and then protein expression was induced with 0.4 mM IPTG at 16 °C for 18 h. The purification procedure was modified from the published protocol of our lab, and the common steps include cell extract preparation, Ni-NTA affinity chromatography (NAC), anion exchange chromatography (AEC) and size exclusion chromatography (SEC) ^22^. In addition, an ammonium sulfate precipitation step (ASP) was performed before NAC to remove the nucleic acids in EcAgo. Specifically, the cell extract was slowly supplemented with saturated ammonium sulfate solution up to the final saturation of 55%. Then, the cell extract was incubated on ice for 1 h and the proteins were collected by centrifugation at 13000 g for 40 min. The proteins were resuspended by 10 mL of lysis buffer (20 mM HEPES pH 7.5, 20 mM imidazole, 500 mM NaCl) and subjected to NAC and the following purification steps as described previously ^22^.

#### Size exclusion chromatography – multi-angle light scattering

SEC-MALS (Size Exclusion Chromatograpy with Multi-Angle Light Scattering) was used to analyze the oligomer size of EcbAgaN. Specifically, after purified with SEC, EcbAgaN was loaded onto a Superdex 200 10/300 GL column (Cytiva, Marlborough, MA, USA) pre-equilibrated with 20 mM Tris-HCl pH 7.5, 250 mM NaCl, at 0.5 mL/min flow rate. Then, Wyatt Dawn Heleos II detector (Wyatt Technology, Santa Barbara, CA, USA) collected the static light scattering, while the absorbance at 280 nm was monitored by AKTA pure 25 UV detector (Cytiva, Marlborough, MA, USA). Data were collected and analyzed in ASTRA 7 software (Wyatt Technology, Santa Barbara, CA, USA). BSA monomer is used as a known molecular weight standard.

#### Nuclease assay

The oligos used as substrates in the nuclease assay are listed in Table S4. The substrates genomic DNA and pUC19 plasmid were extracted from *E. coli* DH5a, while the PCR product substrate was generated by PCR amplification using pUC19 as template and PXS79 and PXS80 primers (Table S3). To analyze the substrate specificity of EcbAgaN, 100 nM FAM-labeled single strand (ss) DNA (OXS1), double strand (ds) DNA (OXS1/OXS2), ssRNA(OXS3), dsRNA (OXS3/OXS4) and DNA/RNA (OXS5/OXS6 or OXS7/OXS8) duplexes were incubated with 200 nM EcbAgaN in the presence of 20 mM Tris-HCl (pH 7.5), 5 mM MgCl_2_ and 5 mM MnCl_2_ at 37 °C for 10 min. Then, the reactions were analyzed by denaturing polyacrylamide gel electrophoresis, and the gel was imaged by a Typhoon 5 laser-scan (Cytiva, Marlborough, MA, USA). The oligos used as the substrates or to prepare the substrates by annealing are listed in Table S4. The same reaction mixtures containing dsDNA (OXS1/OXS2) and ssDNA (OXS1) as substrates were also used to analyze the activity of EcbAgaN mutants. In the assays, the monomer concentration of EcbAgaN was calculated.

To analyze the metal dependency of EcbAgaN, 100 nM FAM-labeled dsDNA (OXS1/OXS2) was incubated with 200 nM EcbAgaN in the presence of 20 mM Tris-HCl (pH 7.5) as well as indicated metal ions or EDTA at 37 °C for 10 min. The reactions were analyzed as described above.

To analyze whether EcbAgaN degrades plasmid and genomic DNA, ∼200 ng pUC19, pCDF-EcAgo^743^, pET28T-EcbAgaN, and *E. coli* genomic DNA were incubated with a gradient of EcbAgaN (0, 50, 100, 200 nM) at 37 °C for 40 min, respectively. Then, the samples were separated by agarose gel electrophoresis, stained with EtBr, and imaged using Molecular Imager Gel Doc EX system (NewBio Industry, Tianjin, China).

To analyze the possible nuclease activity of EcAgo, 100 nM labeled ssDNA (OXS1) and dsDNA (OXS1/OXS2) substrates were incubated with 500 nM EcAgo^731^, with or without 100 nM unlabeled 5P-RNA (OXS13) and target DNA (OXS16) at 37 °C for 1 h. Then, the reactions were treated with 2 mg/mL Protease K (Thermo Fisher Scientific, Waltham, MA, USA) in the presence of 5 mM CaCl_2_ at 55 °C for 30 min. At last, the samples were analyzed by denaturing polyacrylamide gel electrophoresis as described above.

To analyze whether EcbAgaN is activated by EcAgo upon guide and targeting binding, 500 nM EcAgo^731^ was firstly incubated with 500 nM 5P-RNA (OXS13) guide at 37 °C for 15 min, followed by a subsequent incubation with 500 nM target ssDNA (OXS16), nontarget ssDNA (OXS17), target ssRNA (OXS18) and nontarget ssRNA (OXS19) at 37 °C for 15 min respectively. Then, ∼200 ng genomic DNA and 40 nM EcbAgaN were supplemented into the reaction mixtures, which were then incubated at 37 °C for 1 h. Mock incubation using the protein storage buffer and water was performed as controls. At last, the samples were treated with 2 mg/mL Protease K and analyzed by agarose gel electrophoresis as describe above. The plasmid degradation and PCR product degradation assay was performed in the same procedure as the genomic DNA degradation assay except that ∼200 ng pUC19 plasmid DNA and PCR product was treated with 100 nM and 80 EcbAgaN, respectively.

#### Electrophoretic mobility shift assay (EMSA)

To analyze the nucleic acid binding properties of EcAgo, 100 nM of four different 3’-FAM labeled nucleic acid substrates (5P-DNA (OXS10), 5OH-DNA (OXS9), 5P-RNA (OXS12) and 5OH-RNA (OXS11)) were incubated with a gradient of EcAgo^731^ (125, 250, 500, 1000 nM), respectively, in a 10 μL mixture containing 20 μM Tris-HCl pH 7.5, 5 mM MgCl_2_ and 5 mM MnCl_2_ at 37 °C for 15 min. After incubation, the reaction samples were supplemented with 4 μL loading dye containing 50% glycerol, 0.1% bromophenol blue and 0.1% xylene cyanol, and loaded onto 8% native polyacrylamide gels. The electrophoresis was performed in 0.5×TB buffer (44.5 mM Tris, 44.5 mM boric acid) at 100 V for 1 h. At last, the fluorescent signal was visualized using Amersham Typhoon 5 (Cytiva, Marlborough, MA, USA).

To analyze the target binding specificity of EcAgo-guide complex, 500 nM EcAgo^731^ was incubated with 100 nM unlabeled 5P-RNA (OXS13) at 37 °C for 15 min. Then, aliquots of the mixture were diluted to 125 nM and 250 nM EcAgo^731^, respectively. After that, the mixtures were incubated with 100 nM of FAM-labeled target DNA (OXS5), nontarget DNA (OXS14), target RNA (OXS15) and dsDNA (OXS1/OXS2) at 37 °C for 20 min, respectively. The reactions were also analyzed by native polyacrylamide gel electrophoresis as described above.

#### Cytotoxicity assay

In general, the cytotoxicity of the long-B pAgo systems was analyzed in two ways, i.e. protein expression was induced when the cells were grown on plates or in liquid medium, hereafter, the plate induction assay and the liquid medium induction assay.

For the plate induction assay, single colonies of the transformants containing individual long-B pAgos and the associated proteins, the long-B pAgo systems and/or the trigger plasmids were grown in LB medium containing corresponding antibiotics at 37 °C overnight. The cultures (0.1 mL) were transformed into fresh medium (10 mL) containing corresponding antibiotics and grown at 37 °C for ∼3 h. Then, the cells were serially diluted and dropped onto LB agar plates containing the corresponding antibiotics and inducers. After the plates were incubated at 37 °C for 16 h, the numbers of the colonies were counted and colony formation units per mL (CFU/mL) were calculated.

For the liquid medium induction assay, single colonies of the transformants containing individual long-B pAgos and the associated proteins, the long-B pAgo systems and/or the trigger plasmids were grown in LB liquid medium containing corresponding antibiotics overnight. The cultures were transformed into fresh medium at a ratio of 1:100 with corresponding antibiotics (Figure 6C-D, Figure S7) or without them (Figure 3A-C). The cultures were grown for ∼60 min up to an OD_600_ of ∼0.25, when the inducers, including arabinose, aTc and/or IPTG as indicated in the figure legends, were supplemented. Then, aliquots of the cultures were sampled at indicated time points to analyze cell viability and plasmid maintenance, genomic DNA integrity, NAD^+^ level, membrane polarity and permeability as indicated in figure legends. Cell viability and plasmid maintenance were analyzed by measuring CFU/mL on antibiotic-free plates and the plates containing corresponding antibiotics, respectively. The OD_600_ of the starting cultures for the growth curve analysis for BPAS system and BPAM system was adjusted to 0.1.

#### Genomic DNA extraction and analysis

The cells expressing wild type EcBPAN system with pCDF-EcAgo^743^ and pET28T-EcbAgaN, or the mutant systems using pET28T-EcbAgaN-M1 and pET28T-EcbAgaN-M2 instead, or containing the corresponding empty vectors were transformed from overnight cultures into 100 mL antibiotic-free LB medium. At an OD_600_ of ∼0.25, protein expression was induced by 80 ng/mL aTc and 50 μM IPTG. At 0, 1, 2, 4 h post induction (hpi), cells from 2 mL of the cultures were collected by centrifugation. Genomic DNA was extracted using HiPure Bacterial DNA Kit (Magen, Guangzhou, China) following the manufacturer’s instruction, and at the final step, the genomic DNA was eluted by 30 μL water. Then, 1 μL of the DNA samples were loaded on a 0.7% agarose gel, run for 45 min at 150 V in 1x TBE buffer and stained with EtBr. The results were imaged and analyzed using Molecular Imager Gel Doc EX system (NewBio Industry, Tianjin, China).

#### Flow cytometry analysis of DNA content distributions

The cultures were prepared as described in the **Genomic DNA extraction and analysis** section, and the samples for flow cytometry analysis were prepared following the protocol established in our group with some modifications ^22^. Specifically, 300 μL of the cultures were mixed with 700 μL absolute ethanol at 2 hpi and incubated at 4 °C overnight. Before stained, the cells were collected by centrifuging at 5000 rpm for 5 min and washed with 1 mL 1x PBS buffer. The cells were collected by centrifugation again and resuspended in 30 μL 1x PBS buffer supplemented with 2 mg/mL DAPI (Thermo Scientific, Waltham, MA, USA), and stained for at least 1 h on ice in darkness. Then, the cell suspensions were diluted to a final volume of 1 mL by 1x PBS buffer and loaded onto a cytoflex-LX flow cytometer (Beckman Coulter, Brea, CA, USA) with a 375 nm laser, and a dataset of at least 40,000 cells was recorded for each sample. For each cell, the values of fluorescence signal at 450 nm (DAPI signal), FSC (forward scattered light), and SSC (side scattered light) were measured. The results were analyzed and visualized by FlowJo v.10.8.1 (BD Biosciences, Franklin Lakes, NJ, USA).

#### Quantification of cellular NAD^+^ level

The cells expressing EabAgaS or its mutant, or EaBPAS system or the mutant system in the presence of pCDF-EGFP, or containing pBAD24 as empty vector were grown in corresponding antibiotics as described in the liquid medium induction assay. At an OD_600_ of ∼0.25, protein expression was induced by 0.2% arabinose. At 2 hpi, the OD_600_ of cultures were normalized to ∼0.4, and the cells from 1 mL of the cultures were collected by centrifugation and washed by 1 mL 1x PBS buffer. The cells were subjected to the measurements of NAD^+^ level using the Coenzyme I NAD(H) Content Assay Kit (Solarbio, Beijing, China) following the instructions of the manufacturer. Specifically, the cells were resuspended by 500 μL of the acid extraction buffer (Solarbio, Beijing, China), and lysed by the sonicator bath (amp 20%, 2s ON/2s OFF, 2 min duration) at 4 °C. The cell extracts were centrifuged to remove cell debris at 12000 g for 10 min, and the NAD^+^ level of supernatant was quantified by the MTT (Methyl Thiazolyl Tetrazolium) assay following the instructions. Finally, the OD at 570 nm of each sample (OD^sample^) and the corresponding control (OD^sample_control^), where the NAD^+^ in the sample was neutralized before supplementation of MTT, was measured. The relative NAD^+^ content of each sample was calculated using the following equation:

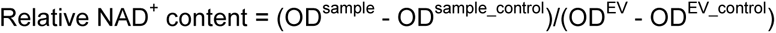

Five biological replicates were performed for the NAD^+^ level assay. EV: empty vector.

#### Flow cytometry analysis of membrane polarity and membrane permeability

The cells containing pBAD24-EaAgo-EabAgaM, pBAD24-EaAgo-EabAgaM_ΔTM or pBAD24 as empty vector were grown in the presence of 100 μg/mL ampicillin as described in the liquid medium induction assay, and the protein expression was induced by 0.2% arabinose. At 2 hpi, ∼10^6^ cells were collected from the cultures by centrifugation (the cell number is calculated based on the estimation that 1 mL of OD_600_=1 culture contains 10^9^ cells). Then, the cells were resuspended in 50 μL 1x PBS buffer containing 1 μg/mL DiBAC_4_ (Sigma-Aldrich, St. Louis, MO, USA) for membrane polarity analysis, or in 50 μL 1xPBS buffer containing 1 μL of dye mix containing SYTO9 and PI in the ratio 1:1(LIVE/DEAD BacLight bacterial viability kit, Thermo Scientific, Waltham, MA, USA) for membrane permeability analysis. Then, the fluorescence signal at 525 nm of the DiBAC_4_-stained cells was collected by the cytoflex-LX flow cytometer, while the fluorescence signal at 525 nm (SYTO9) and 610 nm (PI) was collected for the membrane permeability analysis. All of the data was analysed using FlowJo v.10.8.1 (BD Biosciences, Franklin Lakes, NJ, USA).

#### Purification of EcAgo^743^ for nucleic acid extraction

The cells containing pBAD24-EcAgo^743^ and pCDF-EGFP (+pCDF), pBAD24-EcAgo^743^– EcbAgaN (+bAgaN), pBAD24-EcAgo^743^–EcbAgaN and pCDF-EGFP (+pCDF, +bAgaN) were grown in LB medium containing corresponding antibiotics at 37 °C up to an OD_600_ of ∼0.7. Protein expression was induced by 0.2% arabinose for EcAgo^743^ and 80 ng/mL aTc for EcbAgaN at 37 °C for 4 h. For the cells containing pCDF-EGFP, 400 μM IPTG was also supplemented to induce the T7 promoter. Then, the cells were collected by centrifugation, resuspended by lysis buffer (20 mM HEPES pH 7.5, 20 mM imidazole, 500 mM NaCl) and disrupted by French press. The cell extracts were subjected to NAC purification and the proteins were eluted by a gradient of imidazole. The eluted proteins were analyzed by SDS-PAGE to confirm the expression of EcAgo^743^. Meanwhile, the cells from 10 mL of the same culture were also collected and used for extraction of total RNA, respectively.

#### Extraction of EcAgo^743^–copurified nucleic acids

To extract the copurified nucleic acids, 500 μL of the protein solution was treated with 200 μg/mL Protease K for 1 h and then supplemented with 500 μL phenol/chloroform/isoamyl alcohol (pH 8.0, 25:24:1), followed by a brief vortexing. The sample was centrifuged at 16000 g for 20 min at 4 °C. The upper phase (about 400 μL) was transferred into a new tube and mixed with 40 μL 3 M NaAc (pH 5.2) and 500 μL isopropanol. After incubated at −20 °C for 1 h, The sample was centrifuged at 16000 g for 20 min at 4 °C. The pellet was washed with 1 mL pre-cooled 70% ethanol and dried for 30 min at room temperature. Finally, the nucleic acids in the pellets were resuspended in 50 μL DEPC water.

#### Labeling and treatment of the EcAgo^743^–copurified nucleic acids

One μL of the nucleic acids was treated with FastAP Thermosensitive Alkaline Phosphatase (Thermo Fisher Scientific, Waltham, MA, USA) in a 10-μL mixture containing 1 μL of 10X Buffer (Thermo Fisher Scientific) at 37 °C for 30 min. Mock treatment using water instead of FastAP was performed as controls (-FastAP). Then, the samples were heated at 90 °C for 10 min. One μL of the heated samples was labeled with [γ^32^P]-ATP (PerkinElmer, Waltham, MA, United States) by T4 polynucleotide kinase (PNK, Thermo Scientific) using the forward reaction buffer at 37 °C for 20 min. Then, the samples were analyzed by denaturing polyacrylamide gel electrophoresis. The gel was exposed to a phosphor screen and imaged by a Typhoon 5 laser-scan (Cytiva, Marlborough, MA, USA).

The labeled nucleic acids were further treated with RNase A (DNase and protease-free, Thermo Scientific, EN0531) or DNase I (RNase-free, Thermo Scientific, EN0521) for 1 h at 37 °C, and analyzed by denaturing polyacrylamide gel electrophoresis and autoradiography as described above.

#### RNA sequencing and analysis

Total RNA was used to generate sequencing libraries for the transcriptome analysis with NEBNext Ultra RNA Library Prep Kit for Illumina (NEB, USA, Catalog #: E7530L). The library quality was assessed on the Agilent 5400 system (Agilent, USA). The qualified libraries were pooled and sequenced by Illumina NovaSeq6000 sequencing with PE150 strategy (paired-end reads and 150 bp read length) in Novogene Bioinformatics Technology Co., Ltd (Beijing, China).

Small RNA sequencing libraries were generated using NEBNext® Multiplex Small RNALibrary Prep Set for Illumina® (Set 1) (NEB, USA, Catalog #: E7300S). Subsequently, library quality was assessed on the Agilent 5400 system (Agilent, USA). The Qualified libraries were pooled and sequenced on by Illumina NovaSeq6000 sequencing with SE50 strategy (single-end reads and 50 bp read length) in Novogene Bioinformatics Technology Co., Ltd (Beijing, China).

We used Fastp (version 0.23.1) ^53^ to process the raw reads with default parameters. The processed paired-end reads of the transcriptome sequencing were aligned to the genome of *E. coli* BL21 (GenBank: CP053602.1) and to the expression plasmids (pCDF-EGFP and/or pBAD24-EcAgo^743^ or pBAD24-EcAgo^743^–EcbAgaN) using HISAT2 v2.1.0 (default parameters) ^54^. The single-end reads of the small RNA sequencing were aligned to the above genomes and expression plasmids using bowtie v1.3.1 ^55^, with the -v parameter limiting the number of mismatched bases to 1 and other parameters as default. The length distribution of small RNAs was analyzed by samtools stats ^56^. Nucleotide frequency distributions were visualized using the R package ‘ggseqlogo’ after intercepting mapped reads from 1-15 nt using seqkit ^57^. FeatureCounts ^58^ was used to assign reads to genomic features.

#### Phylogenetic analysis

##### Gene neighborhood (operons) analysis and protein sequences

The accession numbers of 192 long-B pAgos were obtained from the previous study ^8^, and the protein sequences were downloaded from Genbank using Batch Entrez (https://www.ncbi.nlm.nih.gov/sites/batchentrez?). To analyze the associated proteins of long-B pAgos, the neighborhood containing 10 genes from both upstream and downstream of long-B *pAgo* genes was analyzed manually. This reveals that the og_15, og_44 and og_100 genes are invariably organized in the same operon with long-B *pAgos* with conserved operon structure (Figure 1E), while *og_54* is usually located between the 2^th^ and 8^th^ genes upstream of long-B *pAgos*. Then, if the upstream or downstream genes do not encode any protein from the four ogs, we analyzed whether the long-B *pAgos* form operon-like structures with their adjacent genes. If not, the long-B pAgos are considered to have no associated proteins (marked as none in Figure 1A), while the long-B *pAgos* that form operon-like structures with their adjacent genes are considered to have associated proteins other than the four ogs (marked as other in Figure 1A). The protein sequences of og_15, og_44 and og_100 members were also downloaded from Genbank using Batch Entrez.

To select the long-B systems that can be characterized using *E. coli* as a proper host, the protein sequences of og_15, og_44 and og_100 members from the previous study ^8^ were used as queries to search their homologues in the non-redundant protein sequences database with the NCBI blastp suite. Then, we selected the BPAN system and BPAS system from Gammaproteobacteria, and the BPAM system from *Elizabethkingia anophelis*, a mesophilic pathogen (Figure 1E and Table S1). The gene clusters and neighborhoods of the systems were confirmed manually.

##### Phylogeny construction

Homologous sequences were aligned with MAFFT using the automated strategy (v7.490) ^59^. Phylogenetic trees were constructed using maximum-likelihood method by FastTree. The results were saved as newick files and imported into iTOL to plot unrooted tree (v6.5.8; itol.embl.de) ^60^.

#### Tanglegram

The phylogenetic trees of long-B pAgos, and og_15, og_44 or og_100 proteins were imported into R environment (v4.2.1). The tanglegrams were visualized using the cophyloplot function of ‘ape’ R package (v5.6-2) ^61^ with the association information between long-B pAgos and their respective associated proteins.

#### Bioinformatics analysis

The structures of long-B pAgo-associated proteins were predicted by AlpfaFold2 ^62^ using the COSMIC^2^ platform (https://cosmic2.sdsc.edu/). Protein sequence alignment of EcbAgaN homologues and GbbAgaS homologues was performed using Clustal W, and the results were visualized with ESPript (https://espript.ibcp.fr/ESPript/ESPript/). Transmembrane region was predicted with DeepTMHMM (https://dtu.biolib.com/DeepTMHMM).

### QUANTIFICATION AND STATISTICAL ANALYSIS

The cell viability assay, NAD^+^ level assay and growth curve assay were performed in 3-5 biological replicates as indicated in the figure legends, of which the average values are shown in the graphs. The statistical analyses were performed with Excel. The unpaired t-test was used to calculate the p-value: <0.05 = *; <0.01 = **, <0.001=***.

The correlation of the assigned small RNA sequences and transcriptome RNA sequences was analyzed using Origin 2018, with the Pearson correlation coefficients and corresponding p-values calculated.

