## Supplementary Figures for "Long-B prokaryotic Argonaute systems employ various effectors to confer immunity via abortive infection"

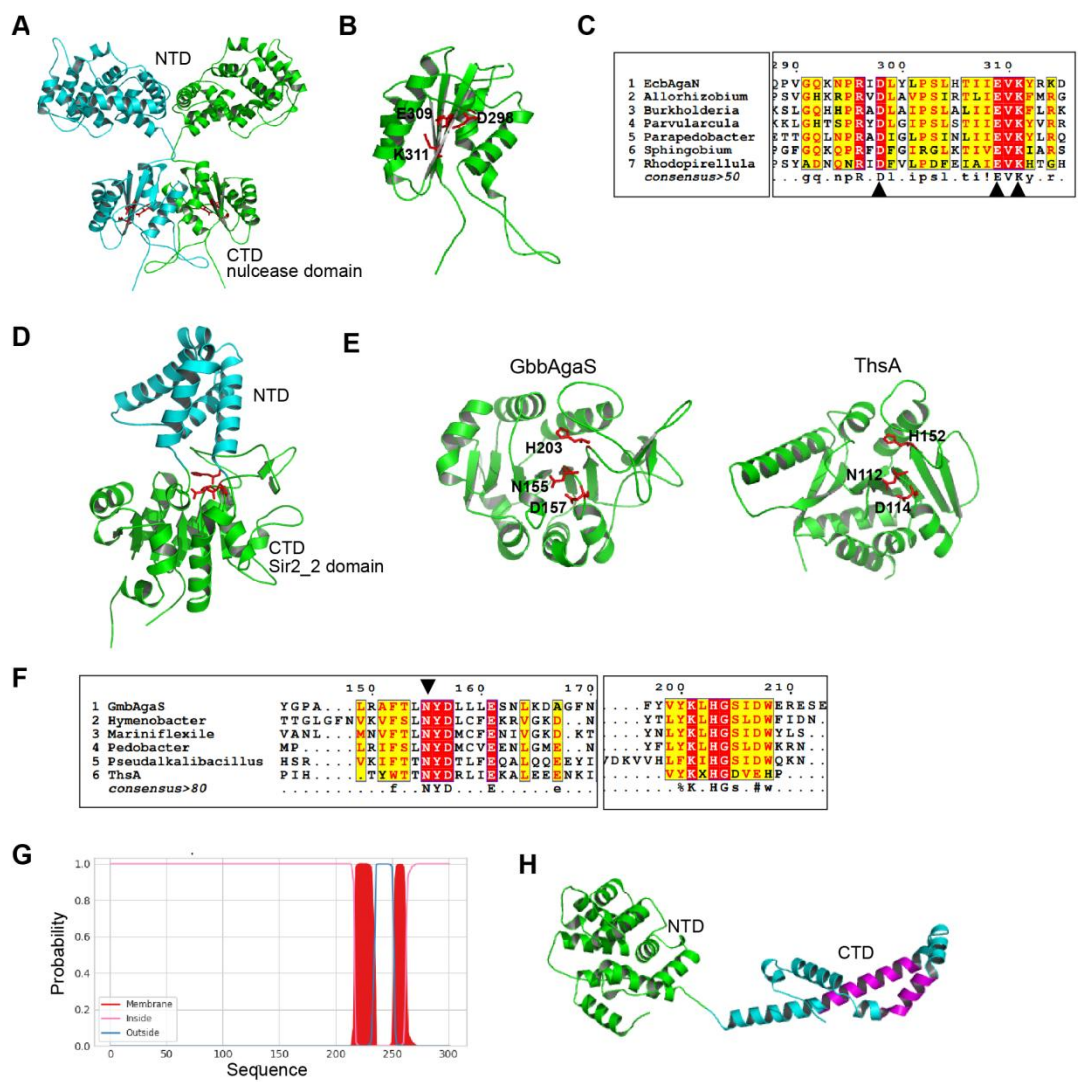

Figure S1. Bioinformatic analysis of long-B pAgo-associated proteins. Related to Figure 1, 2, 6 and 7.

A. Structural prediction of the EcbAgaN dimer by AlphaFold2. The N-terminal domain (NTD) and the C-terminal domain (CTD, the nuclease domain) are indicated

B. A close-up view of the nuclease domain of EcbAgaN. The predicted catalytic sites are indicated.

C. Sequence alignment of EcbAgaN and other representative og\_15 members with the genus name of the source organism shown. Only the conserved region containing the catalytic sites is shown. The arrows indicate the residues for mutagenesis.

D. Structural prediction of GbbAgaS by AlphaFold2. The CTD is the Sir2\_2 domain.

E. Structural comparison of the Sir2\_2 domain of GbbAgaS and the Sir2 domain of ThsA (PDB: 6LHX). The catalytic sites are shown.

F. Sequence alignment of GbbAgaS and other representative og\_44 members, as well as ThsA. Only the conserved region containing the catalytic sites is shown. The arrow indicates the residue for mutagenesis.

G. Transmembrane prediction of EabAgaM by DeepTMHMM.

H. Structural prediction of EabAgaM by AlphaFold2. The predicted transmembrane region is shown in magenta.

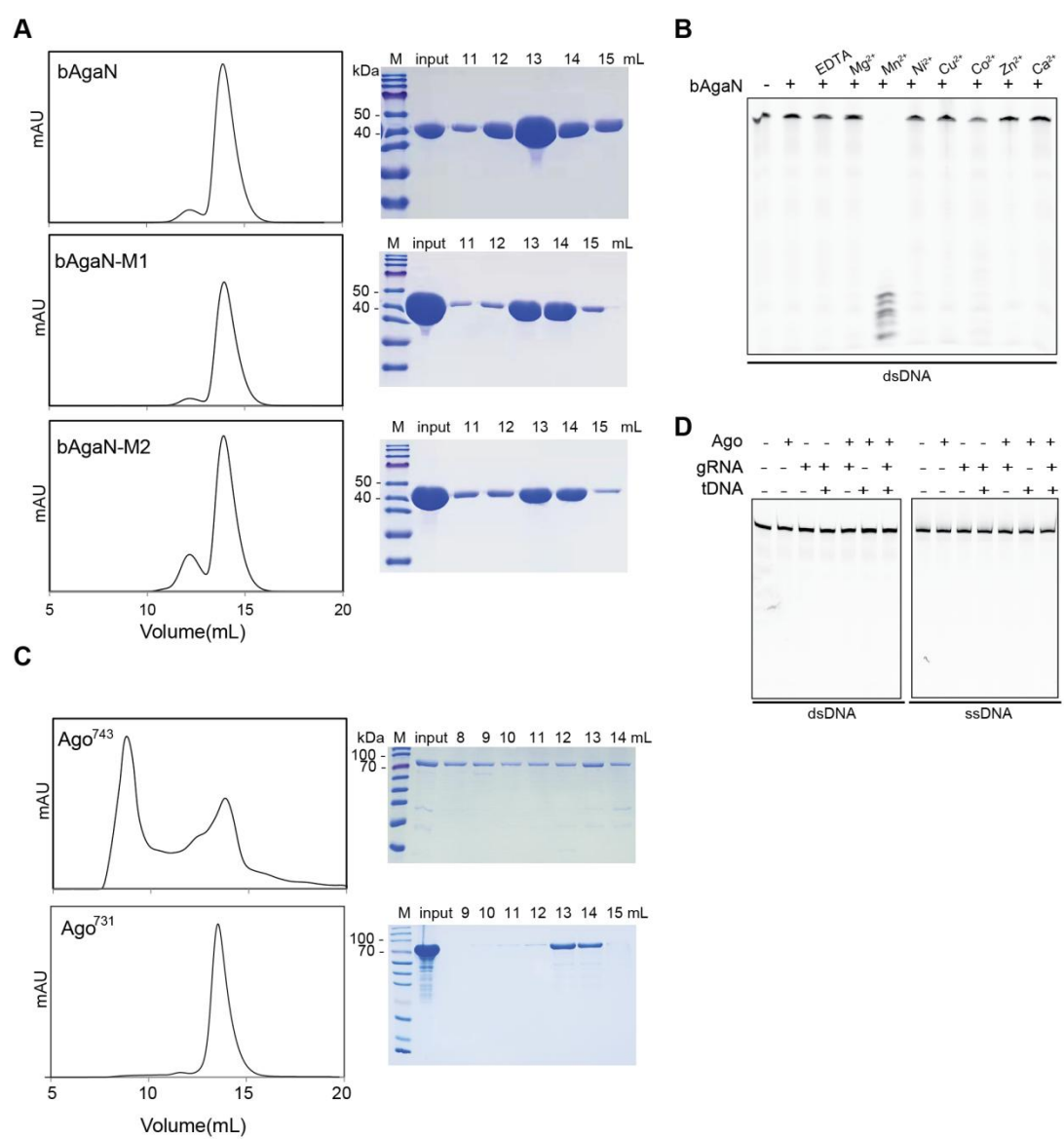

Figure S2. Protein purification and in vitro characterization of EcBPAN system. Related to Figure 2 and 5.

A. Gel filtration profiles of purified EcbAgaN and its mutants (left panels), and SDS-PAGE analysis of the gel filtration samples (right panels). Input: the samples that were loaded onto the gel filtration column. M: protein marker.

B. Metal-dependency of EcbAgaN. FAM-labeled dsDNA was incubated with EcbAgaN in the presence of EDTA or indicated metal ions, and then analyzed by denaturing polyacrylamide gel electrophoresis.

C. Gel filtration profiles of purified EcAgo<sup>743</sup> and EcAgo<sup>731</sup> (left panels), and SDS-PAGE analysis of the gel filtration samples (right panels).

D. EcAgo does not cleave target ssDNA or dsDNA. FAM-labeled target ssDNA or dsDNA was incubated with EcAgo<sup>731</sup>. Guide RNA and/or non-labeled target ssDNA were also supplemented in aliquots of the reaction. Then, the samples were analyzed by denaturing polyacrylamide gel electrophoresis.

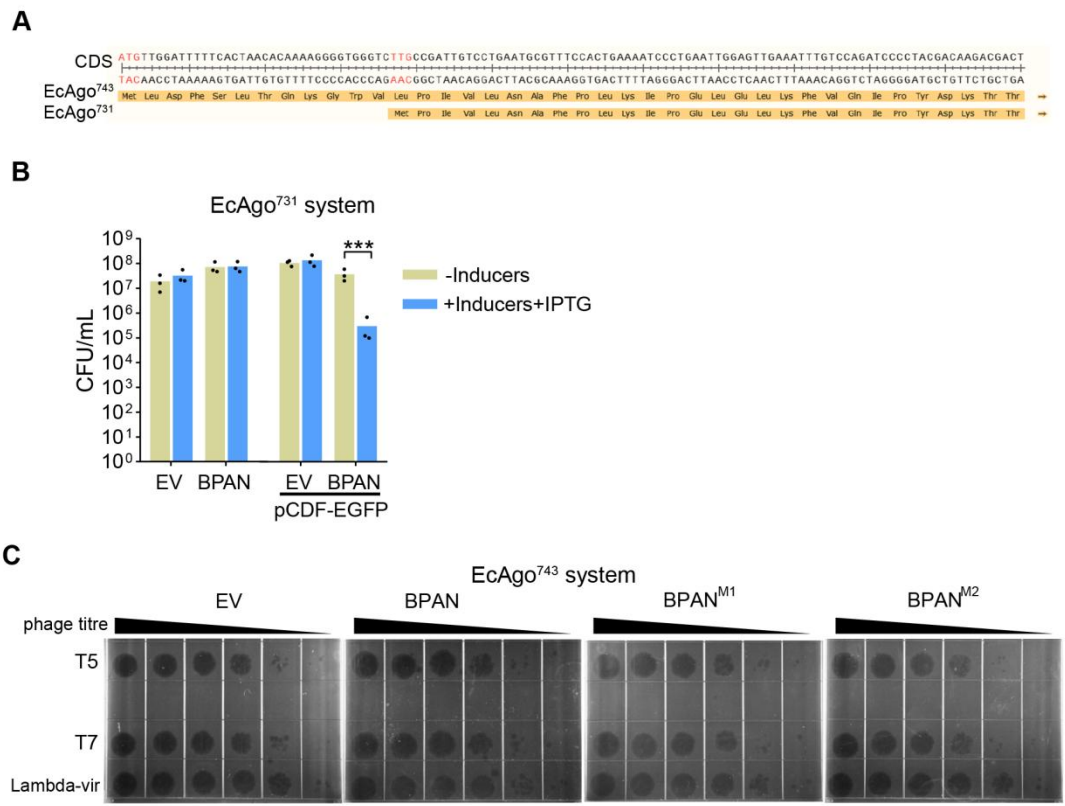

Figure S3. In vivo characterization of EcBPAN system. Related to Figure 3.

A. Two predicted starting codons of *EcAgo*. The coding sequence (CDS) and amino acid sequences of *EcAgo*<sup>743</sup> and *EcAgo*<sup>731</sup> respectively are shown. The two predicted starting codons are marked in red.

B. The *EcAgo*<sup>731</sup>-expressing systems was activated by pCDF-EGFP. Cells containing empty vector (EV) or the *EcAgo*<sup>731</sup>-expressing EcBPAN system in the presence or absence of pCDF-EGFP were plated onto the plates with or without inducers (arabinose and aTc) and IPTG. CFU/mL was calculated and the average of three biological replicates are shown, with individual data points overlaid. \*\*\*: $p < 0.001$.

C. EcBPAN system does not confer immunity against selected phages. The phages were serially diluted and dropped the bacterial lawns expressing EcBPAN system or the mutated systems.

EV: empty vector.

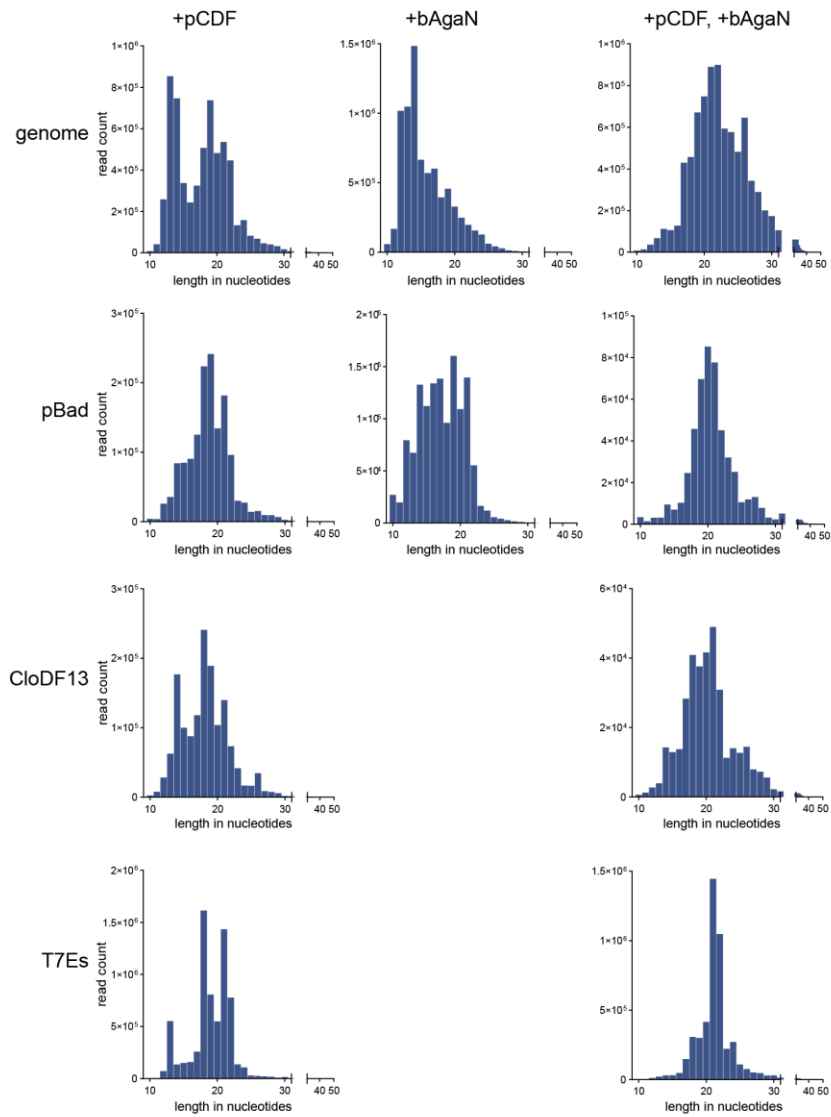

Figure S4. Length distribution of the EcAgo-associated small RNAs assigned to genome, pBAD plasmids, and the CloDF13 origin and the T7 expression cassettes (T7Es) of the pCDF-EGFP plasmid. pBAD can be pBad24-EcAgo<sup>743</sup> or pBad24-EcAgo<sup>743</sup>-EcbAgaN, depending on the samples. Related to Figure 4.

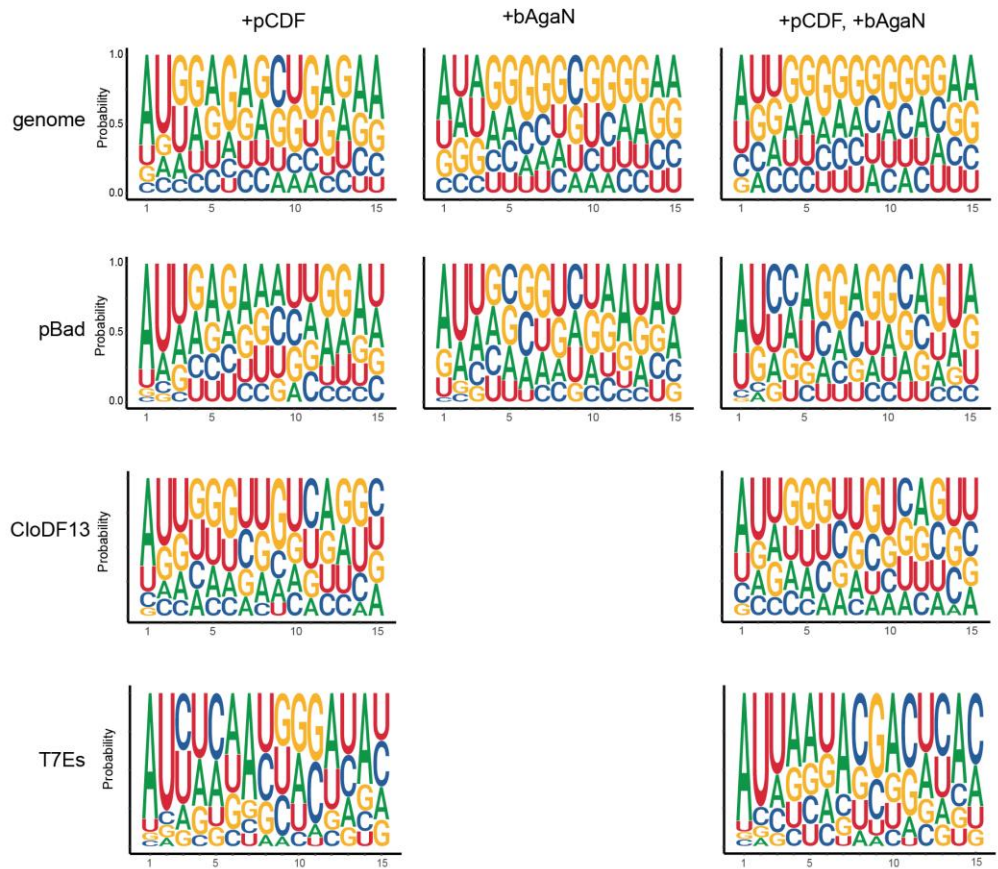

Figure S5. Nucleotide bias of the EcAgo-associated small RNAs assigned to genome, pBAD plasmids, and the CloDF13 origin and the T7 expression cassettes (T7Es) of the pCDF-EGFP plasmid. pBAD can be pBad24-EcAgo<sup>743</sup> or pBad24-EcAgo<sup>743</sup>-EcbAgaN, depending on the samples. Related to Figure 4.

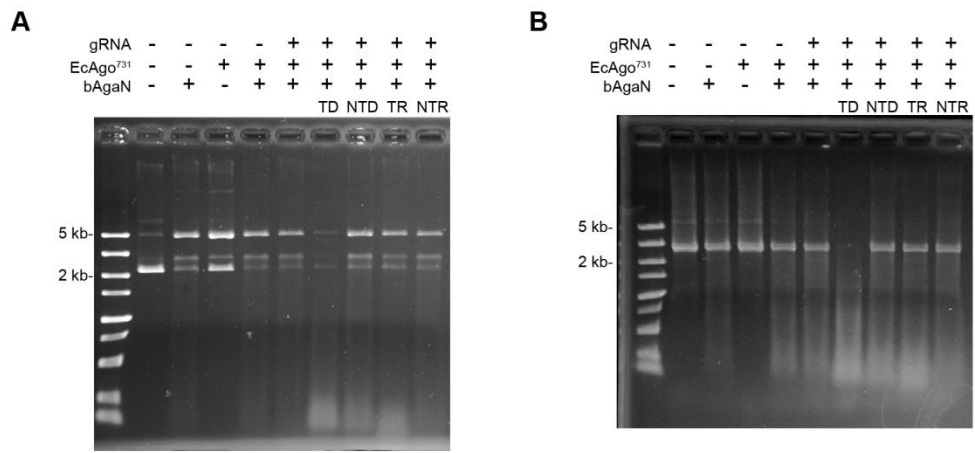

Figure S6. Degradation of plasmid (A) and PCR product (B) by EcbAgaN with or without activation. Related to Figure 5.

About 200 ng pUC19 or PCR product was treated with EcbAgaN in the presence of EcAgo<sup>731</sup>, 5P-RNA (gRNA) and/or target ssDNA or other oligonucleotides. TD: target ssDNA; NTD: nontarget ssDNA; TR: target ssRNA; NTR: nontarget ssRNA.

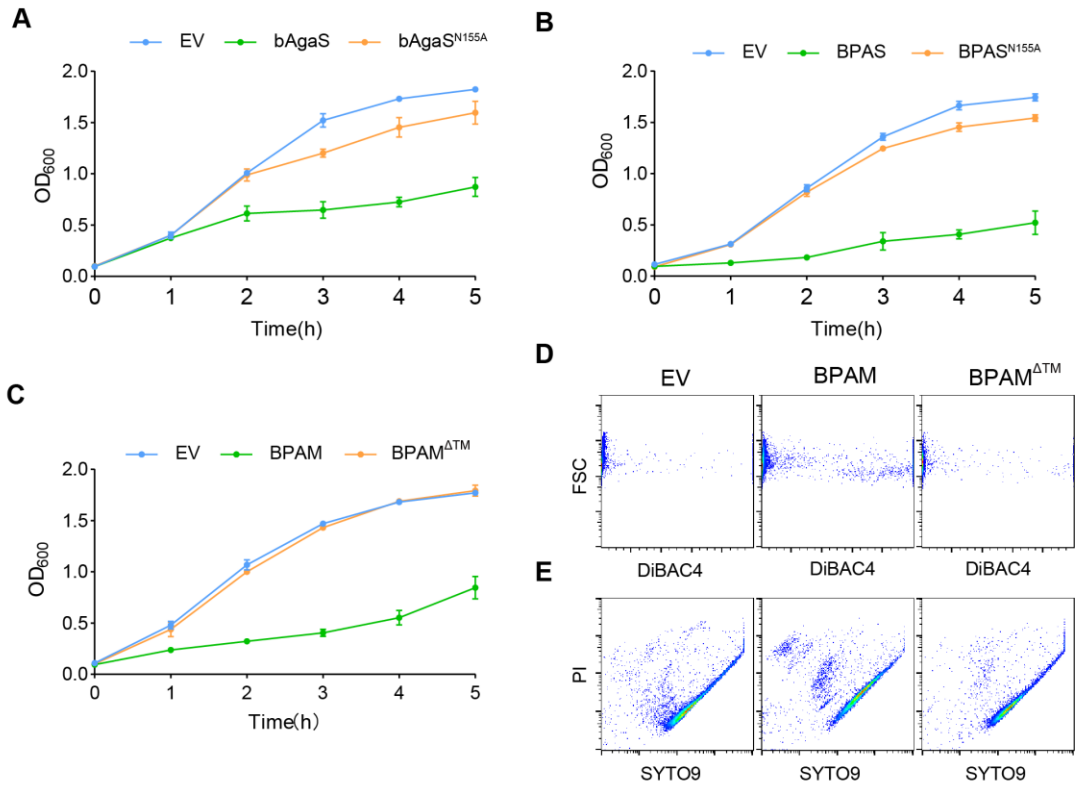

Figure S7. Analysis of the cytotoxicity of the GbBPAS system and the EaBPAM system. Related to Figure 6 and 7.

A. Growth curves of the cultures carrying EV, wild type and mutant GbbAgaS proteins.

B. Growth curves of the cultures carrying EV, wild type and mutant GbBPAS systems in the presence of pCDF-EGFP.

C. Growth curves of the cultures carrying EV, wild type and mutant EaBPAM systems.

E. DiBAC<sub>4</sub> staining of the cells carrying EV, wild type and mutant EaBPAM systems after 2 h's induction.

E. Live/dead (SYTO9/PI) staining of the cells carrying EV, wild type and mutant EaBPAM systems after 2 h's induction.

EV: empty vector.

- Table S1 Proteins analyzed in the study. The names, accession numbers, source strains, and
  sequences are listed. Related to Figure 1-7.
- Table S2 Plasmids used in the study. Related to Figure 2-7.
- Table S3 Primers used in the study. Related to Star Methods.
- Table S4 Oligos used in the study. Related to Figure 2 and 5, Figure S2.
- Table S5 Alignments of sequenced RNA reads. Related to Figure 4.
